# A non-oscillatory, millisecond-scale embedding of brain state provides insight into behavior

**DOI:** 10.1101/2023.06.09.544399

**Authors:** David F. Parks, Aidan M. Schneider, Yifan Xu, Samuel J. Brunwasser, Samuel Funderburk, Danilo Thurber, Tim Blanche, Eva L. Dyer, David Haussler, Keith B. Hengen

## Abstract

Sleep and wake are understood to be slow, long-lasting processes that span the entire brain. Brain states correlate with many neurophysiological changes, yet the most robust and reliable signature of state is enriched in rhythms between 0.1 and 20 Hz. The possibility that the fundamental unit of brain state could be a reliable structure at the scale of milliseconds and microns has not been addressed due to the physical limits associated with oscillation-based definitions. Here, by analyzing high resolution neural activity recorded in 10 anatomically and functionally diverse regions of the murine brain over 24 h, we reveal a mechanistically distinct embedding of state in the brain. Sleep and wake states can be accurately classified from on the order of 10^0^ to 10^1^ ms of neuronal activity sampled from 100 μm of brain tissue. In contrast to canonical rhythms, this embedding persists above 1,000 Hz. This high frequency embedding is robust to substates and rapid events such as sharp wave ripples and cortical ON/OFF states. To ascertain whether such fast and local structure is meaningful, we leveraged our observation that individual circuits intermittently switch states independently of the rest of the brain. Brief state discontinuities in subsets of circuits correspond with brief behavioral discontinuities during both sleep and wake. Our results suggest that the fundamental unit of state in the brain is consistent with the spatial and temporal scale of neuronal computation, and that this resolution can contribute to an understanding of cognition and behavior.

## INTRODUCTION

For nearly a century, stereotyped electrical waves traveling across the surface of the brain have been used to define neural activity patterns correlated with sleep/wake state (Berger, 1929). State-related waves are slow, travel across the isocortex, and are detectable in anatomically distributed structures (Gervasoni et al., 2004; Volgushev et al., 2006). These waves require multiple seconds of observation for identification and can be measured through the scalp, which displays filtered activity averaged across many millimeters of isocortex (Burle et al., 2015). In effect, state-related waves reflect a powerful mechanism of widely distributed electrical coordination, well poised to structure the global state of millions to billions of neurons over seconds to hours. As a result, the neural basis of sleep and wake states is generally understood to be a brain-wide phenomenon and orders of magnitude slower than the submillisecond precision of neural encoding of active behavior (Ding et al., 2016; Lee & Dan, 2012).

However, contemporary studies of brain function and animal behavior are revealing complex state-related dynamics spanning multiple spatiotemporal scales, prompting calls for a reevaluation of traditional perspectives (Nir & de Lecea, 2023). Specifically, there is significant heterogeneity in oscillatory activity within both sleep and wake (Routtenberg et al., 1968; Sainsbury et al, 1987; Harris & Thiele, 2011; Engel et al., 2016; Lacroix et al., 2018; Simor et al., 2020). The low frequency waves that define sleep and wake travel across the isocortex but may be enriched or impoverished in different regions at any moment in time (Huber et al., 2006; Nir et al., 2011; Liu et al., 2022). There is evidence that sleep and wake states may intrude on one another to some extent, even in the course of normal behavior (Emrick et al., 2016; Jang et al., 2022; Soltani et al., 2019; Vyazovskiy et al., 2011; Harris & Thiele, 2011; Engel et al., 2016). The ability of the vertebrate brain to locally regulate states is exemplified by unihemispheric sleep in migratory birds (Rattenborg et al., 2016), marine mammals (Serafetinides & Brooks, 1971), and potentially even humans (Tamaki et al., 2016).

The three broadest states, NREM sleep, REM sleep, and waking, are composed of a hierarchy of substates and nested neurophysiological events, such as sleep spindles and sharp wave ripples. Additionally, the notion of discrete states is somewhat misleading, as transitions between states are characterized by intermediate behavior and neurophysiology (Emrick et al., 2016; Jang et al. 2022; Soltani et al., 2019).

Low frequency (<100 Hz) dynamics are the foundation of previous descriptions of sleep/wake as well as emerging evidence of brief and localized state-related phenomena. These dynamics influence many levels of neuronal activity. For example, delta waves (0.1 - 4 Hz) originating in the prefrontal isocortex drive high amplitude EEG waves that correlate with alternating periods of quiescence and bursting in isocortical neurons (Amzica and Steriade, 1998). Bursting and silence is, unsurprisingly, supported by hyperpolarization and depolarization in neuronal membranes (Volgushev et al., 2006). However, despite the fact that delta waves can be measured in a small patch of membrane, the fundamental unit of the wave is slow (250 ms to 10 s) and widely distributed, coordinating the activity of neurons over multiple centimeters (Volgushev et al., 2006). As a result, local measurements of low frequency waves are understood to reflect global, synchronizing forces (Buzsáki and Schomburg, 2015; Varela et al., 2001; Buzsáki and Draguhn, 2004; Molle et al., 2006). This is the basis of a widely accepted model of brain states in which neuronal activity (fast and local) in distinct circuits is systematically synchronized or desynchronized by oscillations (slow and distributed) (Girardeau & Lopes-Dos-Santos, 2021; Muñoz-Torres et al., 2023).

Due to the physics of waves, it is impossible to consider a substrate of state shorter than the frequency of one cycle (although multiple cycles must be observed for practical purposes). As a result, frequency-based definitions of waves have a minimal resolution that is far slower and larger than the fundamental unit of neuronal activity: the action potential. A wave-based derivation that the fundamental unit of state is slow and global is, while perhaps true, tautological.

We sought to learn the minimally resolvable structure of sleep and wake directly from raw data, independent of assumptions. Convolutional neural networks (CNNs) are well suited to extract the rules of sleep and wake at different spatiotemporal scales. Crucially, CNNs function well with noisy label data - in practice, if a CNN is trained using human labels (based on waves) but learns a more reliable latent signature of state, it can overrule the training label and disagree with high confidence (Rolnick et al., 2018) (Extended Data Fig.8). Using this bottom-up approach, we found that brain states are robustly resolvable in >= 40 ms of data from a single wire placed in any circuit in the brain. Removing oscillatory information below 750 Hz had no impact on accuracy. This suggests that the fundamental unit of brain state is at or below the spatiotemporal order of 10^1^ ms and 10^2^ μm. Fast and local embedding of state provides insights into brain function: individual circuits intermittently switch states independently of the rest of the brain, correlating with brief behaviors in both sleep and wake.

## RESULTS

To empirically evaluate the minimally resolvable fingerprint of brain state in diverse circuitry throughout the mammalian brain, we analyzed a series of long-term, continuous multi-site recordings in freely behaving mice. Briefly, each of nine included animals was implanted with multiple 64-channel microelectrode arrays. To facilitate stable, high signal-to-noise recordings of single unit activity, arrays were composed of tetrodes. Arrays were attached to custom microelectronics in the headstage by flex cables, thus allowing an arbitrary geometry of stereotactic targets. After recovery, recordings (25 KHz) were conducted in the home cage continuously for between 4 and 16 weeks. Preamplification, analog-to-digital conversion, and multiplexing were achieved in the headstage, which was contained in a sparse frame of 3-D printed resin (Fig. 1A). The entire assembly weighed 4 g for a 512 ch (eight module) implant. High resolution video of behavior was collected in parallel. Even in the case of an eight-site implantation, animal movement was similar to unimplanted animals (Fig. S3).

**Fig.1.**
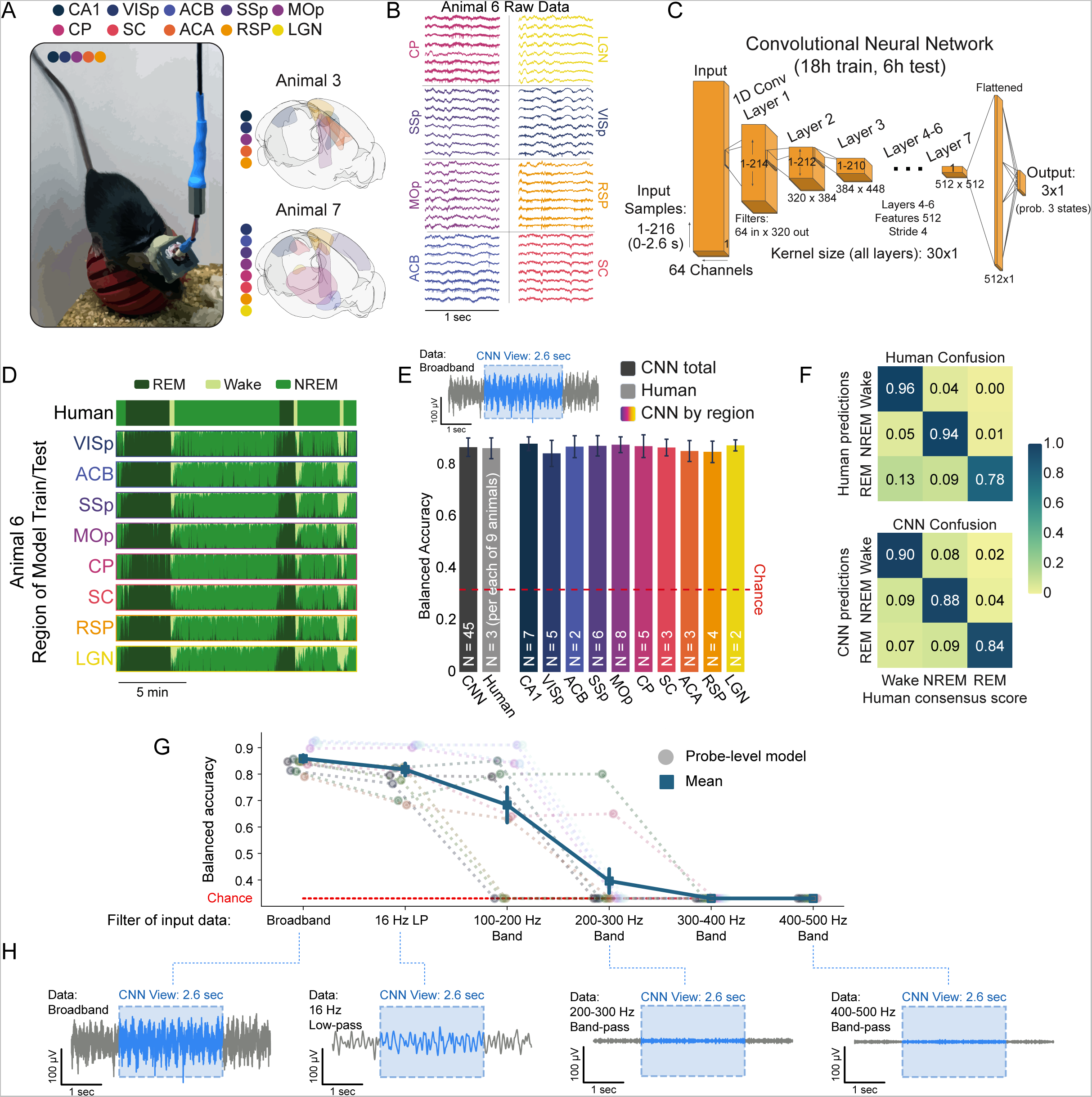
CNNs learn robust signatures of sleep and wake from raw neural data in all brain regions. **A**, Recording protocol. (left) Image of a freely behaving mouse carrying a continuous multi-site recording device in our laboratory. (right) Brainrenders showing examples of implant/recording geometry from two of the nine animals in this study (Claudi et al., 2020). Colored regions indicate recorded circuits. CA1: hippocampus, VISp: primary visual cortex, ACB: nucleus accumbens, SSp: primary somatosensory cortex, MOp: primary motor cortex, CP: caudoputamen, SC: superior colliculus, ACA: anterior cingulate, RSP: retrosplenial cortex, LGN: lateral geniculate nucleus of the thalamus. **B,** 1 s of raw data from 8/64 channels in each of eight implanted brain regions in Animal 6. **C,** The architecture of the convolutional neural network (CNN) used to decode brain state (wake, REM, and NREM sleep) from raw data. Please see the methods section “Convolutional Neural Network Construction and Experiments” for more details. **D,** Human scoring of sleep state (top) versus eight CNNs (bottom) trained and tested independently on eight brain regions in the same animal (y axis of each row represents probability of each of three states at each timepoint). **E,** CNN accuracy relative to a consensus score is slightly but significantly better than individual human experts (left; gray bars, p=0.011, 1-way ANOVA). Colored bars: CNNs trained and tested in each of 10 brain regions show comparable accuracy, text in bars indicates n animals in each region (p=0.180, 1-way ANOVA). Upper inset shows 5 s of raw data (gray) and a 2.6 s sample (blue) that is used by the CNN for classification. **F,** Confusion matrices comparing human scorers (left) and CNNs (right) against consensus scores. Human scorers utilize full polysomnography. CNNs achieve balanced results across three states by observing only raw neural data. **G,** To test the source of state information learned by the CNN, models were trained and tested on filtered raw data from a subset of probes (12 circuits from two animals) : broadband (unaltered raw data), low-pass filtered at 16 Hz, and a series of progressively higher bandpass filters. Balanced accuracy of models is shown as a function of filter. **H,** Visual summary of the various filters applied (gray shows the same 5 s of data in each example). Blue is the 2.6 s window visible to the CNN for scoring.

For inclusion in this study, we selected animals carrying arrays in a minimum of three and a maximum of eight separate brain regions (see Fig. S1 for a map of each animal). Given the diversity of implantation geometries between animals, we ensured that each brain region included in this study was recorded in at least two animals. As a result, we examined a grand total of 10 unique circuits across these recordings (Fig. 1A): CA1 hippocampus (CA1), primary visual cortex (VISp), nucleus accumbens (ACB), primary somatosensory cortex (SSp), primary motor cortex (MOp), caudate/putamen (CP), superior colliculus (SC), anterior cingulate cortex (ACA), retrosplenial cortex (RSP), and lateral geniculate nucleus (LGN). Within each animal’s dataset, we arbitrarily selected a 24 h period for analysis, ensuring only that there was high quality electrophysiological data in every brain region (i.e., evidence of single unit activity on some channels; Fig. S2; Supplemental Table 1). Note that while we used the presence of single unit activity to indicate high quality data, our analyses capitalize on the broadband signal (raw data) unless otherwise indicated (Fig. 1B).

Three human experts independently sleep-scored the 24 h datasets, labeling waking, rapid eye movement (REM), and non-REM (NREM) sleep using polysomnography (Fig. S4). The three experts then met to address points of disagreement in each dataset, thus generating a consensus score that served as the basis of subsequent comparison. Substates within the three states, such as active/quiet wake and sleep spindles, were identified algorithmically and confirmed by human scorers.

### A CNN model can extract sleep and wake states from brief, local observations

Sleep scoring is traditionally conducted taking advantage of EEG, which reflects electrical waves on the dorsal isocortical surface, albeit with low spatiotemporal resolution. In addition, traditional sleep scoring requires some measure of animal motor output, such as EMG. While neural activity throughout the brain is influenced by sleep and wake (Emrick et al., 2016; Jang et al., 2022; Gent et al., 2018; Saper 2006), the degree to which sleep and wake are discernable solely on the basis of these dynamics is less explored. To empirically test this, and allow for circuit-by-circuit variability in state-related dynamics, we trained and tested a unique convolutional neural network (CNN) on the broadband raw data recorded in each circuit (Fig. 1C). Summarily, provided only locally-recorded raw neural data, the CNN attempts to learn rules that consistently predict human-generated labels, even though those labels were derived from EEG and behavior (Fig. 1D).

Each CNN was shown 2.6 s of data at a time (2.6 s CNN) and was trained on 18 h of data and tested on a withheld, contiguous 6 h block of data that spanned a light/dark transition (Fig. S5). Despite the fact that CNNs only observed data from an individual circuit and had no information about animal movement, all three states (REM, NREM, and wake) were robustly separable in 2.6 s increments of data from every circuit. Accuracy was comparable to that of human experts (Fig. 1D-F). Notably, CNNs were effective at detecting brief states, such as microarousals, and in some cases identified examples missed by individual human scorers (Fig. S4F).

CNNs can learn complex patterns from raw data (Rolnick et al., 2018), but understanding the rules learned by models is a widely recognized challenge (Ellis et al., 2022). One approach to this is ablation - by removing key components of a dataset, one can ascertain whether those components are integral to a model’s success. Because human experts rely on low frequency patterns (0.1-16 Hz; Hengen et al., 2016), we hypothesized that CNNs were using the same information. To test this, we applied a series of band-pass filters to data from each circuit in two animals (a total of 12 circuits) - after filtering, only a specific range of frequencies remained. In each circuit, a new series of 2.6 s CNNs was trained and tested on each of five band-passed datasets: A) 0.1 - 16 Hz, B) 100 - 200 Hz, C) 200 - 300 Hz, D) 300 - 400 Hz, and E) 400 - 500 Hz (Fig. 1G,H). Canonical state information is richly embedded in the 0.1 - 16 Hz band, but gamma power (30 - 100 Hz) changes by state as well. If CNNs required canonical oscillatory information, we expected accuracy to decline across these filters. Consistent with this, the 0.1 - 16 Hz model performed within 3% of broadband models (Fig. 1G). CNNs were progressively impaired by the remaining band-passes: all models reached chance levels by 300 - 400 Hz. Taken together, these data demonstrate that CNNs can robustly identify REM, NREM, and wake in nothing but 2.6 s of raw neural data from any circuit and that, consistent with nearly a century of observation (Berger, 1929), models rely on low frequency information to achieve this (Fig. 1H).

### Sleep/wake structures circuit activity at the kilohertz and 10^1^ ms scale

The lowest band-pass suggests that complete information about states is available at frequencies below 16 Hz, and the progressive band-passes confirm that state information declines to 0 by ∼ 200 Hz. However, these results do not rule out the possibility that a mechanistically distinct source of state embedding might emerge in ultra-fast frequencies. We sought to rule this out by training 1 s CNNs on datasets subjected to a series of increasing high-pass filters. In other words, we eliminated low frequency information progressively, allowing all faster information through at each step. Consistent with our prior results, high-passing at 0.3 Hz did not affect model accuracy. Similarly, high-passing above the delta band (4 Hz) had no impact on accuracy. Unexpectedly, this trend continued across almost 4 orders of magnitude (10^-1^ to 10^3^): models maintained full accuracy at 750 Hz, and dropped to 0 only at > 3,000 Hz (Fig. 2A). Models failed in a stepwise fashion: accuracy generally remained above 70% prior to dropping to chance (Fig. S6A). The ∼1 kHz range is understood to contain only fast neuronal events such as action potentials (Chung et al., 2017). Suggesting action potentials as an underlying mechanism, we observed that extracellular action potentials disappeared between 1,000 and 5,000 Hz (Fig. 2A).

**Fig.2.**
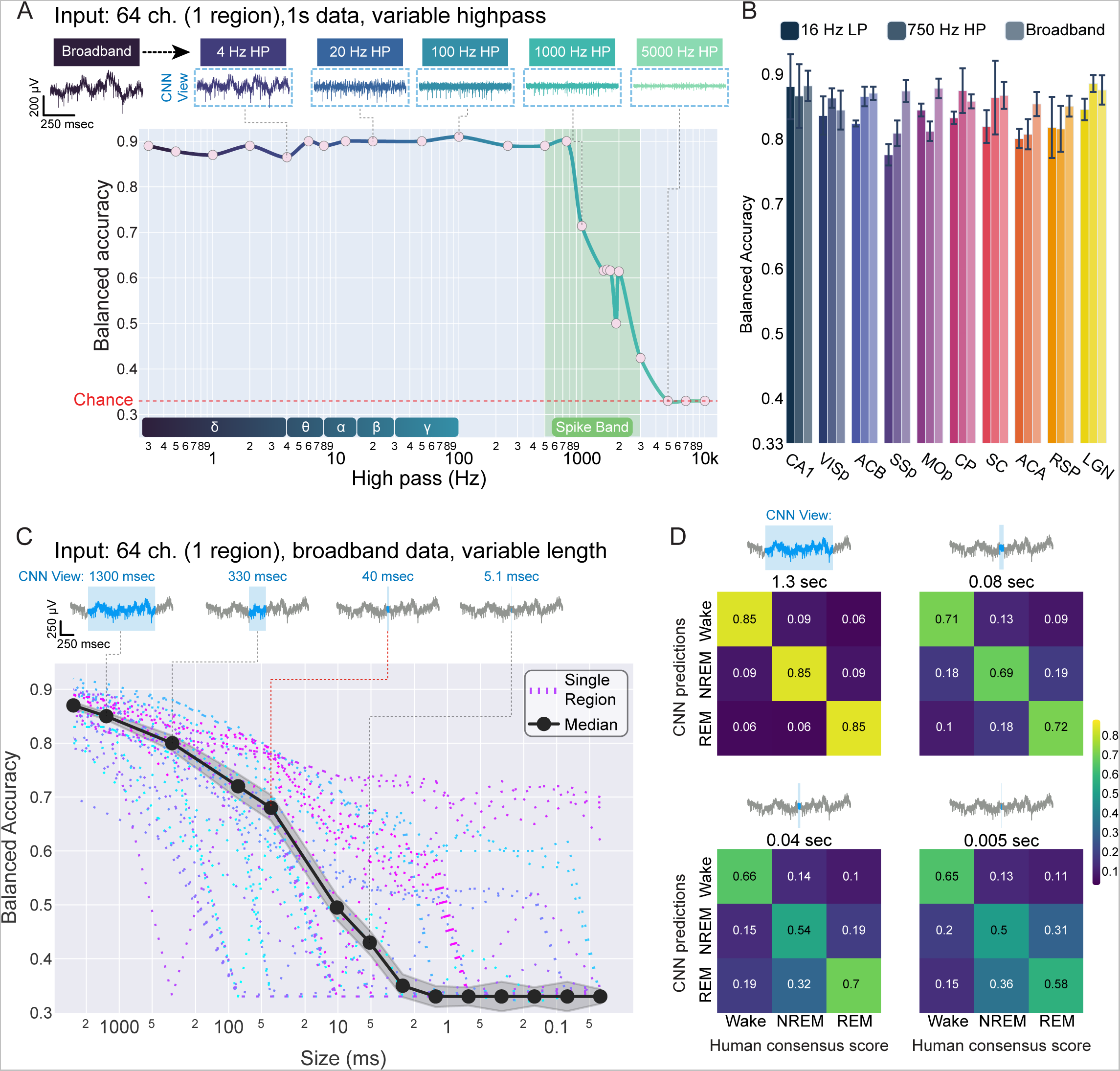
Brain state can be recovered in the kHz band and in only 10^0^ - 10^1^ ms of data. **A**, (top) 1 s of raw neural data subjected to increasing high-pass filters between up to 5,000 Hz. Note that spikes are eliminated between 1,000 and 5,000 Hz. (Bottom) CNN performance as a function of increasing high-pass filters in an example animal. High-pass filtering only decreased brain state information above 1,000 Hz (y = -9.892e-05*x + 0.84, p<2e-16, r2 = 0.43). Lower inset: graphical depiction of classical oscillatory bands used to define states. **B,** Bar plot of summarizing model accuracy by circuit when trained and tested on 16 Hz low-passed data, 750 Hz high-passed data, or broadband (unfiltered) data from all animals. Broadband accuracy was slightly but significantly higher than low-pass accuracy (p = 0.020; linear mixed effects: model_accuracy ∼ filter * circuit + (1 animal) where animal is a random effect) and high-pass accuracy (p=0.014). **C,** (top) 1 s of raw data (gray) overlaid with progressively shorter CNN input sizes (blue). CNNs could only observe an individual sample for each classification. (bottom) Balanced accuracy of CNNs trained and tested on progressively reduced input size. Each recorded circuit (n=45 implants, 9 animals, 10 circuits) is plotted individually (dashed lines), with the overall median performance illustrated (solid black line). **D,** Confusion matrices of mean performance of all models above chance at four input sizes: 1.3 s, 80 ms, 40 ms, and 5 ms. Values and colors represent class balanced accuracy for all models.

Across regions, 1 s CNNs trained on data high-passed at 750 Hz matched the performance of our baseline models (2.6 s broadband CNNs) as well as models restricted to low frequency information (1 s low-pass CNNs) (Fig 2B). Alongside the failure of band-pass models between 200 and 500 Hz, these data strongly suggest that there is a distinct source of brain state embedding in spike-band frequencies. Despite being separated from traditional metrics by multiple orders of magnitude, models restricted to kHz range data suffer no loss in accuracy.

There is a simple explanation by which spike-band information could merely reflect low frequency rhythms present in EEG recordings: periods of action potential bursting and silence are shaped by low frequency field potentials (Vyazovskiy et al., 2011). Put simply, high-pass CNNs may be learning to reconstruct slow information in the timing of fast events. To investigate this possibility, we took advantage of the timescale of oscillatory information: for example, a 1 Hz wave cannot fit in windows smaller than 1 s. We trained another series of CNNs, this time systematically manipulating the size of input data. We progressively reduced the input size from 2.6 s down to a single sample point, or 1/25,000 s (Fig. 2C). Each model was provided with data from all channels within an individual circuit, but limited to a given input size. While median model performance declined monotonically as a function of input size, CNNs maintained accuracy significantly above chance down to 5 ms (0.43 +/-0.02, chance = 0.33). Given only 40 ms of input data, median CNN accuracy was nearly 70% (0.68, +/-0.02, chance = 0.33). Note that chance is precisely 0.33 - CNNs are provided with balanced datasets, and the mode of failure is characterized by universally guessing a single class (e.g., NREM). We excluded models which failed in this respect from statistical comparisons. Interestingly, we observed relatively balanced performance across the three states all the way down to 5 ms (Fig. 2D); in other words, CNNs did not struggle to identify one of the states relative to the other two. Taken with the high-pass results, these data suggest that all three states impose discriminable structure on neural activity at the millisecond timescale.

Finally, we reasoned that CNNs may be learning to reconstruct slow waves in short intervals by virtue of spatial information gleaned by sampling 64 channels. Succinctly, because a wave travels in space, an instantaneous measurement of voltage on multiple electrodes could provide a measure of wavelength. To address this, we trained another series of CNNs exposed to progressively smaller and smaller input sizes. This time, however, we only passed in data from a single channel. While overall accuracy was slightly reduced and more variable in this heavily constrained situation, we observed qualitatively similar results in all single-channel models from all circuits of all animals (Fig. S6B). Many single-channel models performed significantly above chance down to 1 ms.

### Brain states impose millisecond-scale patterning in neuronal activity

We next sought to understand how sleep and wake state are embedded in neuronal activity at high frequencies and in small intervals. Logically, there are three possible mechanisms by which this phenomenon could be explained. First, models could learn that the average instantaneous voltage differs by state. This is the only possible explanation of performing above chance when testing on a single sample point. Second, variance might differ by state. In this case, two or more sample points could provide insight. Each of these explanations is decodable in fast/small samples, but does not necessitate a fast underlying process. Third, neural activity could be patterned at the millisecond timescale. In this case, the sequence of voltage measurements would carry state information and require a fast organizing process.

To test these possibilities, we trained a series of single-channel CNNs (Fig. 3A) on the same datasets in two conditions. One was low-passed at 16 Hz and one was high-passed at 750 Hz (Fig. 3B). In effect, after high- and low- pass filtering, the same sample was passed into corresponding CNNs for labeling. In these parallel high- and low- pass datasets, we trained a series of CNNs on systematically decreased sample intervals. Finally, to test the hypothesis that high-frequency state embedding might reflect patterned information, we trained another parallel set of models but shuffled each sample prior to input (Fig. 3C). To summarize, progressively smaller chunks of data from a single wire were sleep scored by CNNs in four scenarios: intact low-pass, intact high-pass, shuffled-low pass, and shuffled-high pass.

**Fig.3.**
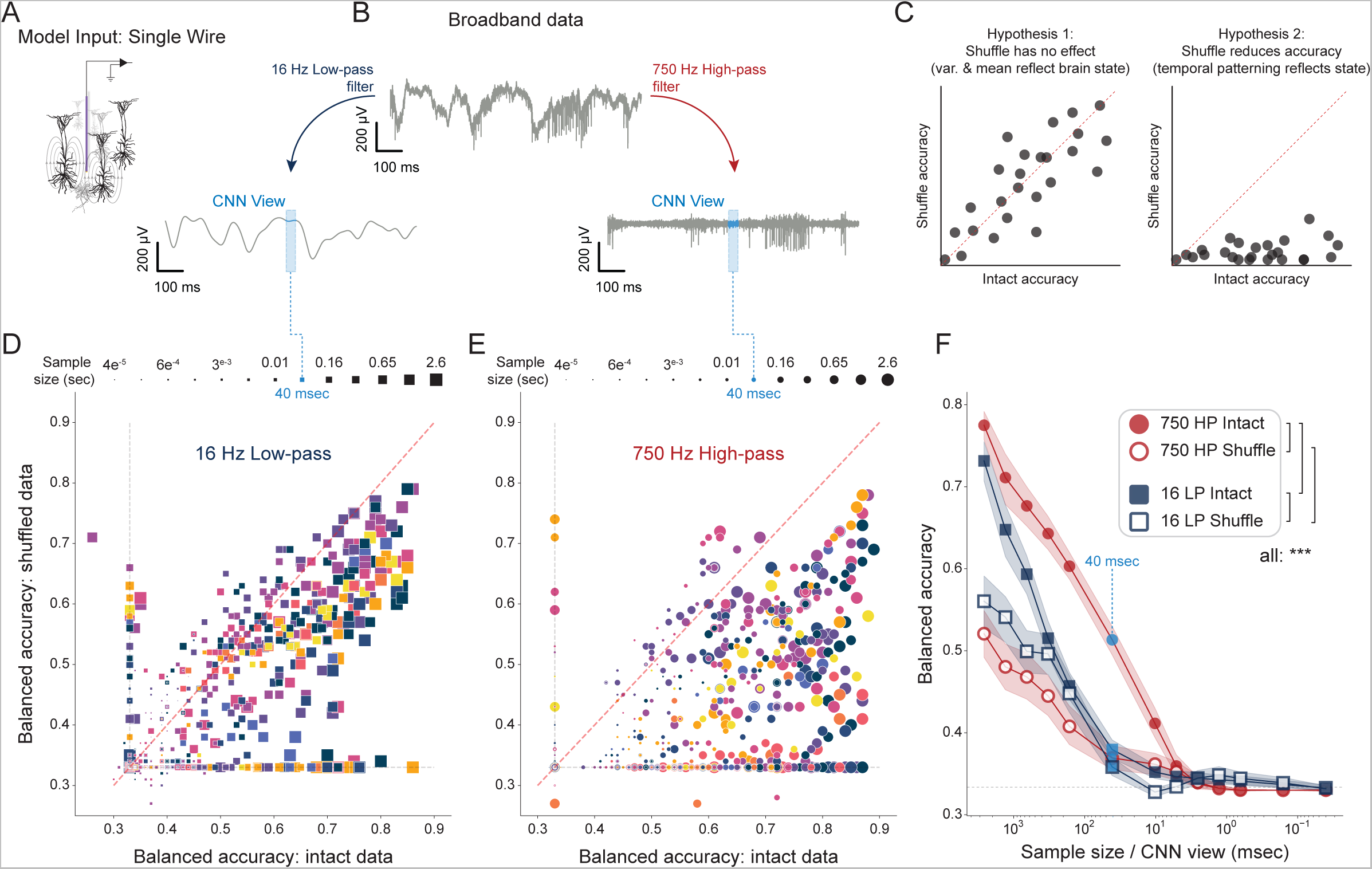
Brain states pattern high frequency neuronal dynamics on the order of 10^0^ to 10^1^ ms. **A**, Experimental overview: model input. Depiction of a single wire (⌀ = 12 μm) placed in a local circuit. Curved lines represent the spatial effects of single neuron current dipoles that influence measured voltage at high frequencies. **B,** Experimental overview: paired low-pass and high-pass models. Parallel models are trained and tested on data from the same single channel (A), one model observing 16 Hz low-passed data, the other observing 750 Hz high-passed data. Blue boxes depict a 40 ms observation interval in each case. **C,** In each condition (low- and high- pass), another pair of models are made: intact and shuffled data (each sample is shuffled prior to training/testing). If shuffling does not reduce the ability of a CNN to decode state (left), state information must be recoverable from sample mean and variance. If temporal pattern is determined by state, shuffling will reduce accuracy (right). **D,E,** Two single channels were selected in each recorded circuit (both with and without high amplitude spiking) for examination in four conditions: high- / low- pass filtering and shuffle / intact comparison. In each condition, models were trained/tested at 13 input sizes (shown along the top) from 2.6 s (65,536 data points) to 0.04 ms (1 data point), yielding a total of 2,028 single-channel CNNs. **D,** 16 Hz low-passed single channel data. Each square shows a pair of intact/shuffled models, the square is colored by the circuit on which they are trained, and the size of the square indicates input size. **E,** The same as D, but for 750 Hz high-passed single channel data. **F,** Summary of models in D and E. Red indicates 750 Hz high-pass, blue indicates 16-Hz low-pass. Filled points are intact data, and open points are shuffled samples. 40 ms is indicated by dashed light-blue line for ease of comparison with other results. *** indicates p < 0.001. Linear mixed effects balanced_accuracy ∼ input size * filter * shuffle + (1|animal) + (1|circuit).

In the time intervals examined (2.6 s - 1/25,000 s), intact low-pass models performed only slightly better than their shuffled counterparts - a comparison of paired intact/shuffled low-pass models reveals a tendency to cluster near the unity line (Fig. 3D). The performance of shuffled and intact low-pass models was only separable down to 655 ms (p <.0001; linear mixed model: balanced_accuracy ∼ sample size * high_low_pass * shuffle + (1|animal) + (1|circuit), where animal and circuit are random effects), after which they converged and then drop to near chance (<40%) by 40 ms (Fig. 3F). These data demonstrate that low frequencies carry some information about brain state at relatively short intervals, but that this information is distributional - the fact that shuffling does not impair performance reveals that models are learning from mean and/or variance.

In contrast, intact 750 Hz high-pass models significantly outperformed their shuffled analogues at all sizes above 2.5 ms. A comparison of paired intact/shuffled models reveals a rightward shift toward intact models (Fig. 3E), consistent with the hypothesis that high frequency/short latency state information is temporally patterned (Fig. 3C). Not only did intact 750 Hz high-pass models outperform their shuffled variants, they also significantly outperformed all other models above 2.5 ms, including intact low-pass models at 2.6 s (p = 0.0036). At 40 ms, the intact highpass model was the only model to perform above 40% (near-chance), yielding an accuracy of 51.3 +/- 0.9%. This is consistent with the results of 64 channel CNNs tested on broadband data (Fig. 2C), which suggested that 40 ms is the minimal sample size that maintains high accuracy. Taken together, these data reveal that high frequency temporal patterning carries state-related information on the timescale of 5 ms and above. Due to the effects of space filtering, it is likely that these patterns are generated within <100 μm of each recording site (Bédard et al., 2006).

Surprisingly, the effect of shuffling was observed similarly on channels both with and without high amplitude spiking. That even non-spiking channels outperformed their shuffled counterparts reveals that the background noise in recordings contains temporally structured information about brain state. The most likely source of this information is low amplitude spiking of nearby neurons, i.e. multiunit-hash (Harris et al., 2016; Trautmann et al., 2019). This may account for the ability of high-pass CNNs to accurately identify state even when input samples contain no obvious single unit spiking.

### Substates and fast neurophysiological events

It has long been recognized that sleep and wake comprise many substates. In addition, stereotyped, intermittent neurophysiological events unfold in a state-dependent fashion. Sharp wave ripples are enriched in NREM sleep and quiet waking (Vanderwolf, 1969; Girardeau et al, 2009), while cortical ON- and OFF- states are primarily associated with NREM sleep (Vyazovskiy et al., 2011). These examples raise two questions about short and fast embedding of brain states. First, could it be that CNNs rely on fast substates and events (e.g., sharp wave ripples) to achieve their accuracy? This is unlikely, because fast events such as sharp wave ripples comprise a tiny fraction of total time in a state. Second, given that key neurophysiological events are defined by rapid and dramatic changes in activity, do such events lead to model errors and confusion?

To address these questions, we used established algorithmic methods to detect sharp wave ripples (Karlsson & Frank, 2009; Kay et al, 2016), sleep spindles (Vallat & Walker, 2021), and cortical ON- and OFF- states (Vyazovskiy et al., 2011) in each of our datasets. In addition, we divided all of waking into two substates, active and quiet wake. We then evaluated the accuracy of 40 ms CNNs in and around these events.

Cortical ON- and OFF- states during NREM did not confuse 40 ms models, despite the fact that individual ON- and OFF- states lasted longer than 40 ms (Fig. 4A,B). This suggests that, despite the fact that the CNN was only able to view data entirely within an ON- (or OFF-) state, there was a reliable latent signature of NREM. Cortical ON- and OFF- states can also occur in waking and REM sleep, albeit less frequently (Vyazovskiy et al., 2011; Funk et al., 2016). To a human observer, 40 ms of data within an OFF-state from REM, NREM and wake are identical, yet 40 ms CNNs correctly identified the superstate of these ON- and OFF- states (Fig. 4C,D). Likewise, the 40 ms CNN was robust to sharp wave ripples (Fig. 4E) and sleep spindles (Fig. 4F). Finally, network activity differences between quiet and active wake did not drive confusion (Fig. S8A,B).

**Fig.4.**
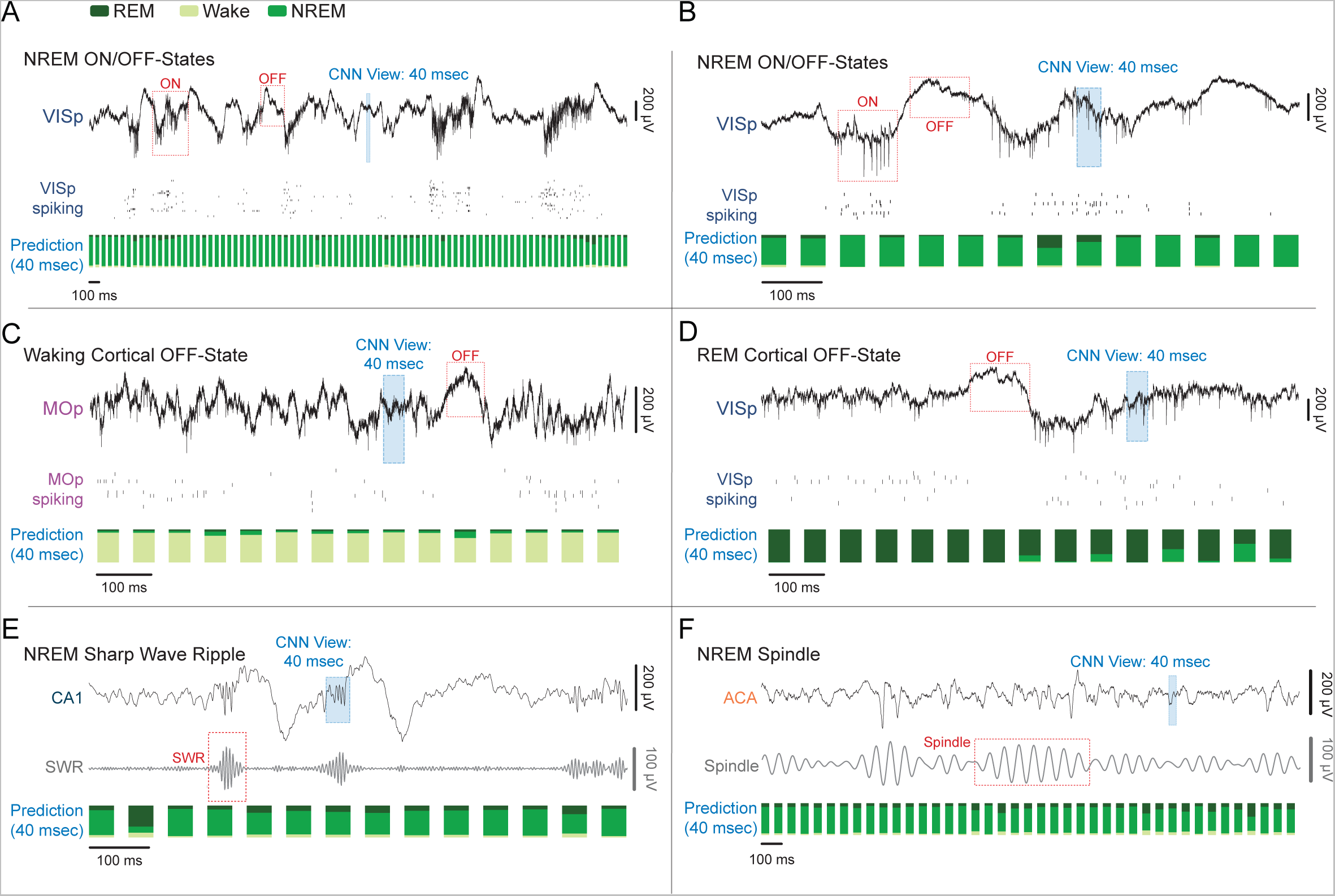
Fast embedding of states is robust to diverse low frequency activity and neurophysiological events. **A**, Top- Broadband trace of several seconds of exemplary high-delta (0.1 - 4 Hz) activity during NREM sleep. Data are recorded in VISp. Red boxes indicate cortical ON and OFF states. Blue box shows the width of an individual input sample used by the 40 ms CNN to predict state. Middle- Raster of subset of VISp single units spiking. Bottom- Stacked barplot of 40 ms CNN prediction probabilities (the three colors in each bar show the probability that the corresponding sample came from each of the three states). To reduce computational burden, the CNN evaluates a 40 ms sample every 1/15 s (hence the slight gaps between samples). **B,** Zoomed 1 s view of ON/OFF-states in VISp. **C,** Example of a waking OFF-state in MOp. **D,** Example of a REM OFF-state in VISp. **E,** Example of a NREM sharp wave ripple (SWR) in CA1 hippocampus. Middle trace shows the same data as top trace but filtered to highlight SWR. **F,** Example of a NREM spindle in ACA. Middle trace shows the same data as top trace but filtered to highlight spindles.

### Microcircuits exhibit brief, independent sleep and wake-like states

We noted that CNNs of all sizes occasionally gave rise to high confidence errors and even disagreed with labels in training datasets (Fig. S7). Interestingly, these disagreements often spanned independent models trained in distinct circuits in the same brain. Given the confidence and independent reproducibility of these events, we reasoned that they may be evidence of two known phenomena. First, micro states have been described extensively (Halasz 1998). Micro arousals are brief (3-15 s) global arousals within NREM sleep (Halasz 1998; Ekstedt et al., 2004) into wake followed by a prompt return to NREM. Microsleeps are equivalent but involve a brief intrusion of sleep into waking. Second, slow waves characteristic of sleep may appear in portions of the cortex of awake animals in conditions of sleep deprivation (Vyzaovskiy et al., 2011; Alfonsa et al., 2023) or inactivity (Andrillon et al., 2021). This is referred to as local sleep.

To account for these, we operationally defined microarousals as brief periods (< 20 s) occurring during otherwise consolidated sleep in which independent models in each circuit identified waking with high confidence (Fig. 5A, left). We applied an equivalent definition to microsleeps, identifying periods during waking when all models briefly identified a sleep state (Fig. 5A, middle). We eliminated these from our analyses. This excluded roughly 10% of nrem-to-wake disagreements and 5% of wake-to-nrem disagreements. After this, we were left with brief states that occurred in only a subset of circuits (i.e., not global), which we termed “flickers” (Fig. 5A, right).

**Fig.5.**
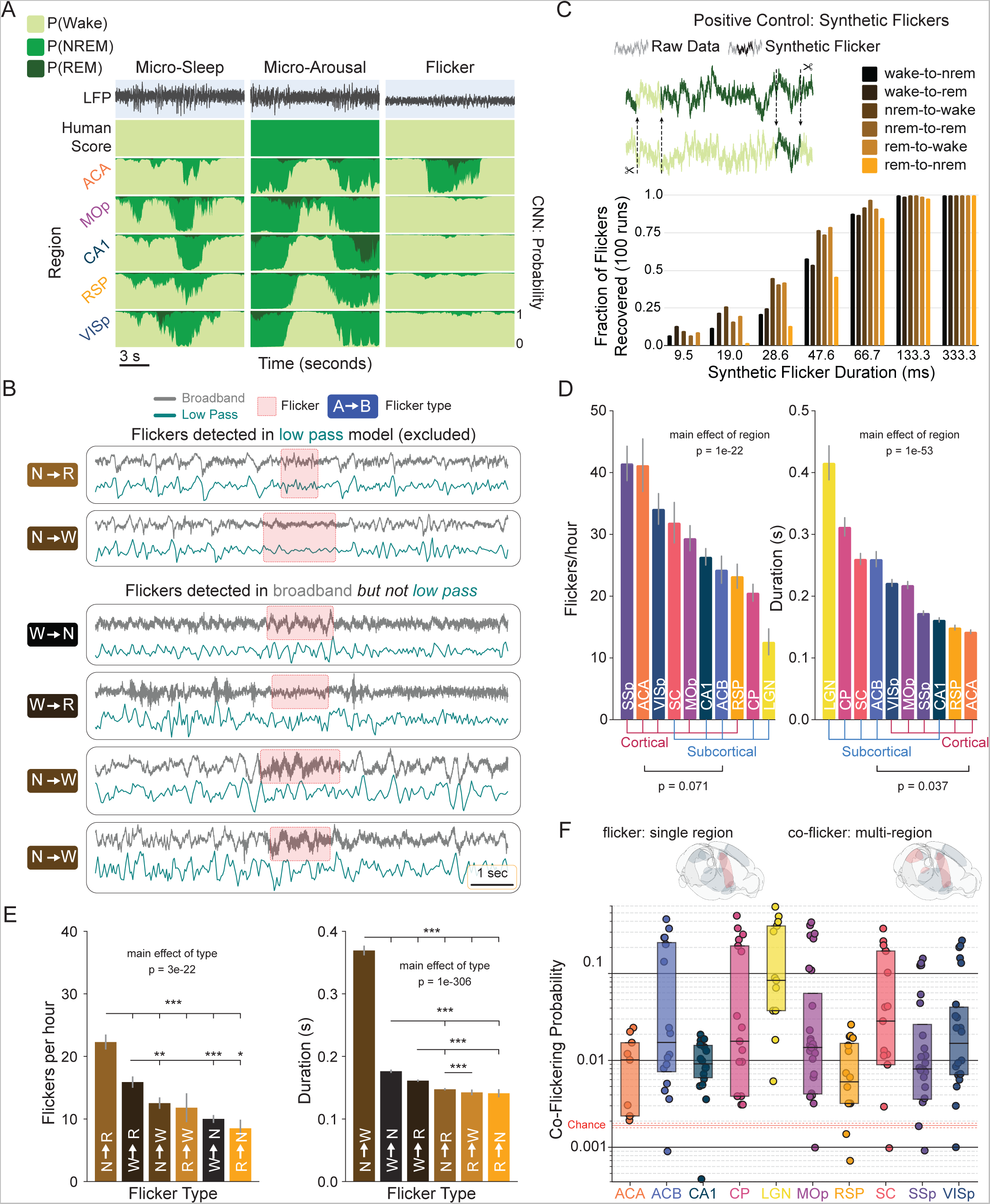
Individual circuits briefly switch states independently of the rest of the brain. **A**, Examples of three forms of disagreement between CNN classification and human consensus scoring of brain state. Top trace is neural broadband. Second row is human scoring of corresponding state. Bottom four rows are outputs of independent CNNs trained in each of four brain regions recorded in the same animal. The left column is a microsleep: all circuits (global) show a brief, high confidence intrusion of sleep into surrounding wake. The center column is a microarousal: all circuits show a brief, high confidence instruction of wake into surrounding sleep. The right column demonstrates a wake-to-NREM “flicker” in the anterior cingulate. Flickers are defined as high-confidence, non-global events that are not detected in low-pass models (B), and are distinct from transitions between states. **B,** Brief and local events were identified in models trained on 16 Hz low-passed data as well as models trained on broadband data. Events detected in both models were excluded from subsequent analyses. (top two rows) Examples of flickers detected in broadband data (gray trace) as well as low-passed data (teal). Red boxes denote the interval identified as a flicker in each model. Flicker type is shown on the left. (bottom four rows) Examples of flickers identified in the broadband but not low-passed data. **C,** (top) Schematic illustrating synthetic flicker positive control. Short segments of data from each state were transposed into segments of each other state. (bottom) Proportion of synthetic flickers captured by CNNs as a function of duration and flicker type. **D,** Flicker rate and duration vary significantly as a function of circuit (linear mixed effects model; Fig. S9A for post-hoc pairwise comparisons). There was a trend towards higher flicker rate in isocortical than subcortical regions (p = 0.071, Spearman Rank Correlation). Subcortical regions exhibited significantly longer flickers than isocortical regions (p = 0.037). **E,** (left) Mean rate of each flicker type per hour. (right) Mean duration by flicker type. See Fig. S9B for post-hoc pairwise comparisons. **F,** Coincident flickers (co-flickers) in two or more anatomically distinct circuits (top illustration, right brainrender) occurred significantly above chance (p<0.001, permutation test). Box plot of co-flickering probability by circuit. Shaded red area between dashed red lines indicates the range of chance levels (min to max) in all circuits. Solid red line is the mean chance level.

We initially hypothesized that flickering reflected local sleep (Vyazovskiy et al., 2011). To evaluate this, we deployed 1 s CNNs on 16 Hz low-passed data sets and identified flickers in this output. Approximately 60% of flickers identified in the broadband model were captured by low-pass models (Fig. 5B). These were consistent with human readable, low frequency local states, such as local slow waves during wake (Vyazovskiy et al., 2011) and REM (Funk et al., 2016), local changes in slow wave activity during NREM (Siclari and Tononi 2017), and local theta power during patterned behavior (Routtenberg et al., 1968; Sainsbury et al., 1986). We eliminated these from our analyses. We also directly evaluated whether flickers represent confusion driven by episodic oscillatory events. Neither ON/OFF states, sleep spindles, nor sharp wave ripples correlated positively with flickers. We reasoned that remaining events might represent transient, local shifts in the high frequency latent patterning of state.

We also note that more than 99% of the excluded global events were also detected by low-pass models. This supports the efficacy of low-frequency monitoring tools (like EEG/LFP) for detection of microarousals and microsleeps.

To test the sensitivity of CNNs to momentary switches of state, we created a positive control. We randomly transposed variable length snippets of data between stretches of REM, NREM and wake (Fig. 5C). 1 s CNNs recovered such synthetic flickers down to 10 ms, and recovered all examples > 66 ms. This suggests that flicker events identified by CNNs are consistent with brief intrusions of one state into another.

Because such flickering appeared to be a plausible neurobiological event, we systematically quantified flicker rate and duration as a function of circuit. Flickers occurred at a rate of 10 - 50 per hour in each circuit (Fig. 5D. p = 1e-22, main effect of circuit, linear mixed model: flicker_rate ∼ circuit + (1 | animal), where animal is a random effect; see Fig. S9A for pairwise comparisons between circuits). Notably, there was a trend for isocortical regions to generate flickers at a higher rate than subcortical regions (p = 0.071, Spearman Rank Correlation). Consistent with our synthetic control (Fig. 5C), flickers were observable down to 67 ms. Mean flicker duration varied significantly by circuit from 142 to 416 ms (Fig. 5D. p = 1e-53, linear mixed model: flicker_duration ∼ region + (1 | animal); see Fig. S9A for pairwise comparisons). Subcortical circuits produced significantly longer flickers than isocortical circuits (p = 0.037, Spearman Rank Correlation).

We observed all six possible flicker types, i.e., combinations of surrounding and flicker states. There was a significant effect of flicker type when comparing rate (Fig. 5E. p = 3e-22, linear mixed model, flicker_rate ∼ type + (1 | animal). See Fig. S9B for pairwise comparisons). The most frequent flicker type was NREM to REM, while the least was REM to NREM. Similarly, there was a significant effect of flicker type on duration (Fig. 5E. p = 1e-306, linear mixed model, flicker_rate ∼ type + (1 | animal). See Fig. S9B for pairwise comparisons). NREM to wake flickers were the longest, while REM to NREM flickers were the shortest.

Interregional coordination is known to play a major role in the emergence and maintenance of sustained arousal states as well as the transient modulation of attentional states (Harris & Thiele, 2011; Poulet & Petersen 2008; Engel et al., 2016; Tan et al., 2014; Zagha et al., 2013; Mitra et al., 2016). We hypothesized that, if flickers represent neurobiologically meaningful substates, their timing might reflect functional connectivity between circuits. In other words, flickering in one circuit might strongly influence the state-related dynamics in downstream circuits. Consistent with this, subnetworks of circuits reliably exhibited coincident flickering far above chance (Fig. 5F). It is interesting to note that the LGN exhibited the highest rate of co-flickering, consistent with the broad connectivity of the thalamus.

### Flickering corresponds to transition-like spiking activity

Our results thus far demonstrate that CNNs identify unique state-related information in the 1 KHz range, which contains action potential traces (Fig. 6A). We hypothesized that single unit spiking might show evidence of state alterations during flickers. If so, this would suggest that flickering corresponds with a meaningful shift in neuronal spiking. Alternately, if unit activity were unaffected by flickering, this would suggest that these brief events detected in the broadband data are statistical anomalies.

**Fig.6.**
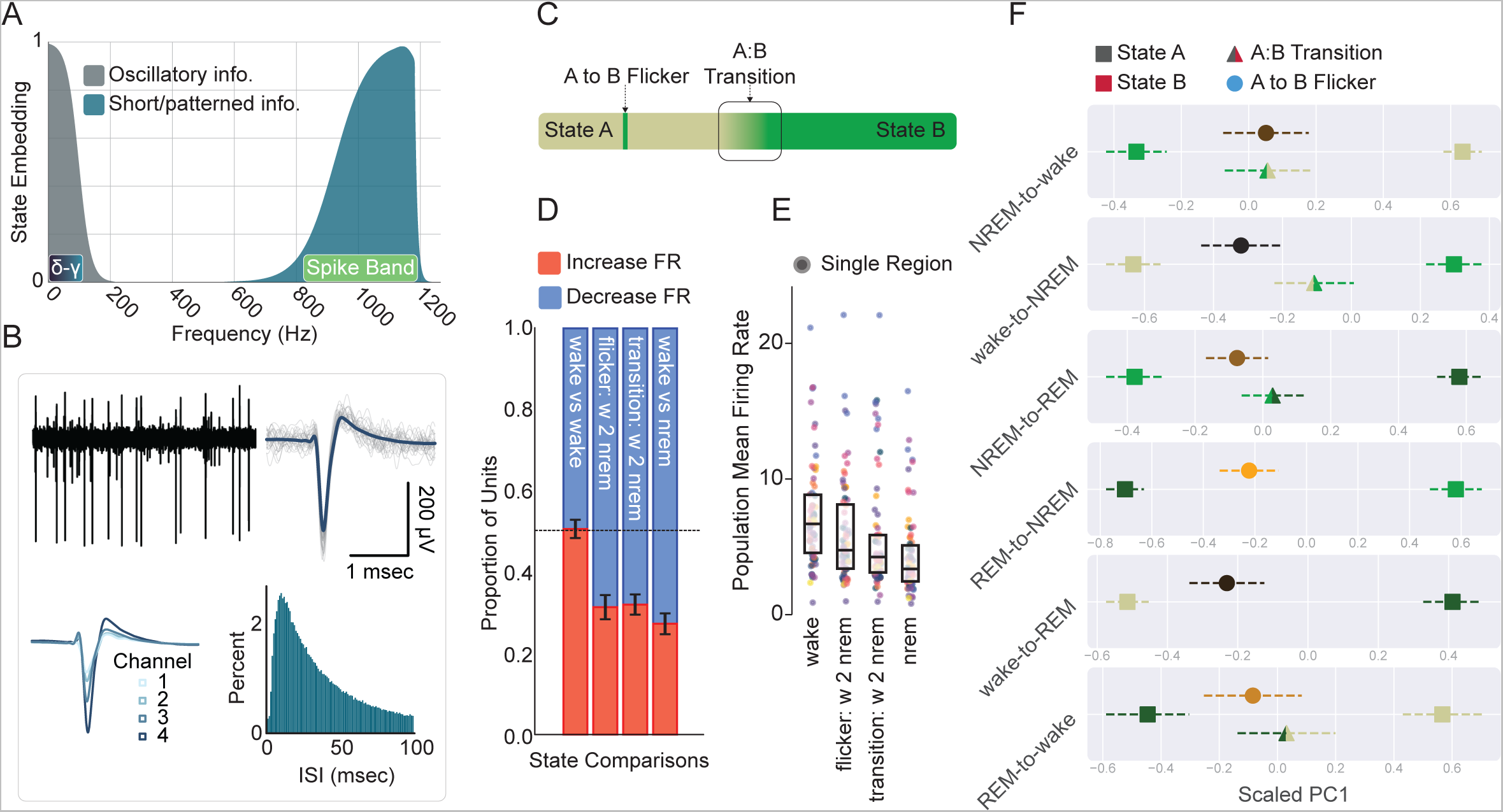
Single neuron spiking shows evidence of flickers detected by CNNs. **A**, Schematic of state information as a function of frequency content. The shaded gray captures state information conveyed by canonical oscillations (δ-γ bar inset). The shaded teal illustrates state embedding in the kHz range, which contains action potential information (spike band). **B,** Example of a single unit. Top left is a broadband trace showing high signal-to-noise spiking. Top right shows the mean waveform (dark blue) of an extracted and spike-sorted single unit (individual traces shown in gray). Bottom left shows the mean waveform across the four channels of a tetrode. Bottom right is a histogram of the unit’s interspike intervals. Note the presence of a refractory period around 0-5 ms. **C,** Conceptual illustration of a flicker and a transition. **D,** The portion of units whose sampled instantaneous firing rate was different relative to a random sample of the surrounding state (wake): wake vs. wake (negative control), wake-to-NREM flicker, wake-to-NREM transition, and wake vs. NREM. See Fig. S10A for all state pairs. Error represents SEM. **E,** The mean single unit firing rate by circuit (color) during wake, a wake-to-NREM flicker, a wake-to-NREM transition, and NREM. See Fig. S10B for all state pairs. Box plots show the intermediate quartiles with outliers as swarm scatter colored by region. **F.** Mean scaled PC1 projections for the six flicker types- surrounding state (A: left square), predicted state (B: right square), A-to-B flicker (triangle), and A-to-B transition (circle) are shown. To incorporate the n animals into estimated variance, error bars are the SEM multiplied by the square-root of n animals. See Supplemental Tables 2-5 for significance of pairwise comparisons based on linear mixed effects model projection by sample type (i.e. flicker, transition, surrounding, predicted).

We spike-sorted all raw data and extracted ensembles of well-isolated single units from every circuit (Fig. 6B and S2). We then separated spiking into bins corresponding to each state, transitions between states, and flickers (Fig. 6C). Many units systematically changed their firing rates as a function of state - the proportion of units displaying alterations between states was maintained both in transitions and in flickers (wake to NREM example shown in Fig. 6D; all combinations shown in S10A). Likewise, population mean firing rates showed consistent shifts between states, in transitions, and in flickers (Fig. 6E, all combinations shown in S10B).

Lastly, we sought to compare the effect of flickers and transitions on individual units whose activity varied substantially between states; it is these units in particular that might be expected to be modulated by flickering. We used principal components analysis (PCA) to examine correlations between the spiking patterns of single units (sets of 100 ISIs) on a circuit-by-circuit basis. By definition, the first principle component (PC1) represented the largest source of variance in unit activity - fortuitously, the axis defined by PC1 reliably separated each pair of states with near perfect accuracy, suggesting that ensemble spiking activity is sufficient to support arousal state classification.

We then projected spiking activity during flickers and transitions onto these axes, allowing us to directly ask whether spiking during flickers showed evidence of a separation from the surrounding state. Ensemble spiking during both flickers and transitions was significantly shifted from the surrounding state towards the flicker state in all six flicker types and all four transition types (Fig. 6F). Transitions were significantly separable from the surrounding state in all regions, and flickers were separable in 9/10 regions (Fig. S10C, Extended Data Table 2-5). In 5/6 flicker types, flicker duration was uncorrelated with position along this axis (the exception was nrem-to-wake flickers, p<0.001). This suggests even very short flickers corresponded to spiking changes. The number of regions a flicker was detected in was also not significantly correlated with its position along this axis. Thus, time periods identified as flickers and transitions by the CNN correspond to significant alterations in single unit activity.

Spiking activity during flickers and state transitions were generally not significantly different; this was the case in 3/4 transition types and 9/10 circuits (Extended Data Table 2-5). K-means clustering was applied along these axes to further explore the relationship between transitions and flickers on a recording-by-recording basis. On each PC1 axis, an optimal k was determined by the silhouette score k = 2.79 +/- 0.10, and flickers co-clustered more frequently with transitions (68.53%) than the surrounding (63.13%) or predicted states (34.14%) (Rousseeuw 1987). This suggests, despite significant variability, the primary configuration along this axis is three clusters: one where the majority is the surrounding state, a second shared by transitions and flickers, and a third populated by the predicted state. Through the lens of single unit firing, flickers are consistent with circuit dynamics during transitions between the surrounding and predicted state.

### Flickering correlates with transient behavioral changes

As a whole, our data suggest that the minimally reliably resolvable unit of brain state is on the order of 10^1^ ms and arises from local circuit activity. One possibility is that this is an epiphenomenon - in other words, resolving states at this scale provides no insight into brain function. Alternatively, understanding states at fast and local resolution could offer novel insight into the structure of natural behavior. Because flickers comprise brief changes in neuronal spiking, we hypothesized that they might correlate with animal behaviors that unfold on a similar timescale.

We used optical flow - the difference between adjacent video frames - as a coarse measure of total activity. Optical flow differed significantly between all pairs of states (Fig. S4G). Interestingly, there was a slight but significant reduction in optical flow during REM relative to NREM, consistent with REM paralysis.

Given that flickers represent momentary discontinuities in state, we next asked if similar discontinuities in behavior (Kramer & McLaughlin, 2001) corresponded with flickering. We algorithmically identified three forms of naturally occurring behavioral discontinuities: 1) motor twitches during extended NREM sleep (Fig. 7A), 2) brief pauses amidst extended locomotor sequences (Fig. 7B), and 3) momentary reductions in slight movements during NREM sleep, such as those associated with muscle tone and respiration (i.e., “freezing”, Fig. 7C). We examined the relationship of these discontinuities to flickering. Curious to know whether a potential relationship might depend on the number of circuits contributing to a flicker, we divided our analyses into single-region flickers and multi-region co-flickers.

**Fig.7.**
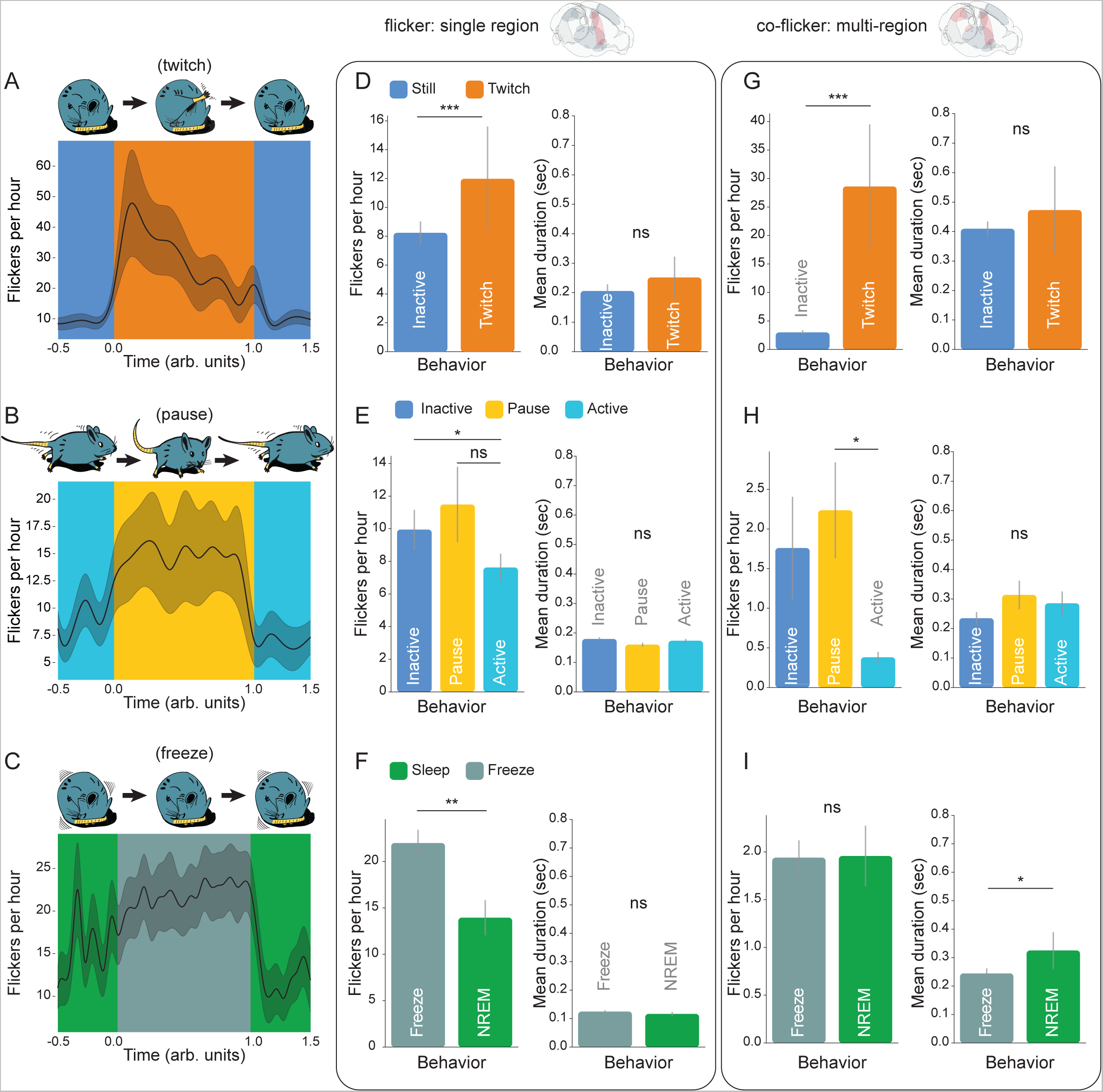
Flickering predicts structure in free behavior. **A**, (top) Illustration of a twitch during inactive NREM sleep. (bottom) NREM-to-wake flicker rate before (dark blue), during (orange), and after (dark blue) twitches. Error shade is SEM. **B,** (top). Illustration of a brief pause during extended locomotion. (bottom) wake-to-NREM flicker rate before (light blue), during (gold), and after (light blue) pauses. Error shade is SEM. **C,** (top) Illustration of brief “freezing” during inactive NREM sleep (i.e., a slight but significant reduction in slight movements associated with muscle tone, respiration, etc.; Fig. S4G). (bottom) NREM-to-REM flicker rate before (green), during (slate), and after (green) freezing. Error shade is SEM. **D,** Rates (left) and durations (right) of NREM-to-wake single-region flickering (top brainrender) as a function of motion states shown in A. **E,** Same as D but for wake-to-NREM flickers during the states shown in B. **F,** Same as D but for NREM-to-REM flickers during the states shown in C. **G,** Rates (left) and durations (right) of NREM-to-wake co-flickers (multi-circuit: top brainrender) as a function of the motion states shown in A. **H,** Same as G but for wake-to-NREM flickers during the states shown in B. **I,** Same as G but for NREM-to-REM flickers during the states shown in C. Error bars for all plots are SEM. * p<0.05, ** p<0.01, *** p<0.001, linear mixed effects: flicker_rate ∼ motor_state * flicker_type * n_circuits + (1 | animal/circuit) or flicker_duration ∼ motor_state * flicker_type * n_circuits + (1 | animal/circuit), n = 45 circuits, 9 mice.

In all three examples of brief behavioral switching, we observed significantly increased flicker rates. Specifically, NREM-to-wake flickers were enriched during sleep twitches (Fig. 7A. p<0.001, linear mixed regression: flicker_rate ∼ motor_state * flicker_type * n_circuits + (1 | animal/circuit), where circuit nested within animal is a random effect), wake-to-NREM flickers were enriched during brief pauses in high activity (Fig. 7B. p = 0.0018), and NREM-to-REM flickers trended towards an increase during freezing in sleep (Fig. 7C. p = 0.0722). In the first two cases (NREM-to-wake and wake-to-NREM), multi-fold increases in co-flickering appeared to drive these effects, suggesting that when more circuits are recruited during a flicker, there is more impact on behavioral structure. Paradoxically, co-flickering masked a significant enrichment of single-circuit NREM-to-REM flickers during freezing in sleep (Fig. 7I. p = 0.006).

These data suggest that short states identified in individual circuits correlate with short behavioral states within sleep and wake. By demonstrating a relationship between flickers and behavior, these data imply that consideration of brain states at the level <100 ms and kHz frequency has the potential to explain meaningful features of brain function and organization.

## DISCUSSION

Sleep and wake states are widely assumed to arise from extended changes in global patterns of brain activity (Ding et al., 2016; Gervasoni et al., 2004; Lee & Dan, 2012; Volgushev et al., 2006). Here we report the unexpected finding that sleep and wake determine distinct, millisecond-scale patterns in microcircuits throughout the brain. We find that individual circuits regularly switch states (which we refer to as “flicker”) independently of other circuits. Flickers demonstrate that fast, state-related patterning of neuronal activity arises locally. Flickers, as brief neurophysiological discontinuities, are enriched in brief behavioral discontinuities in both sleep and wake. This implies that fast, state-related dynamics may contribute to cognition and behavior. Our data suggest a fundamental unit by which state determines brain activity that is distinct from low frequency oscillations.

Our data also suggest that extremely short intervals of neural data contain latent structure that is determined by state. This information takes the form of distinct temporal patterns, although the mechanism behind such a phenomenon remains unclear. An immediate possibility is that of nested oscillations, similar to how 1 Hz slow waves in NREM coordinate sharp wave ripples (150 Hz). However, this is unlikely for three reasons. First, it appears that fast embedding can exist *despite* slow rhythms, a property illustrated by flickering. For example, within NREM slow waves, fast embedding can reveal a flicker (Fig. 5B, bottom two rows. Note that high frequency information is visibly altered during flickers despite ongoing slow wave activity). Second, there is a 500 Hz gap between low frequency rhythms and fast embedding (Fig. 1G,H, and Fig. 6A). Third, analysis of nested events and substates (Fig. 4) reveals that, amidst dramatic changes in low frequency activity, reliable fast embedding is trivially recoverable. As a result, it is not immediately clear how traditional wave information below 100 Hz could establish the kHz frequency fast embedding described here.

However, extensive empirical evidence suggests that slow, broad signals are the foundation of sleep and wake states in the brain (Steriade et al., 1993; Gervasoni et al., 2004). Consistent with this, activation of broadly projecting nuclei in the midbrain and brainstem drives changes in brain state (Carter et al., 2010; Chen et al., 2018; Moruzzi & Magoun, 1949; Li et al., 2022). Our data thus appear paradoxical: sleep and wake can be experimentally controlled by slow and anatomically distributed mechanisms, yet cannot account for millisecond patterns nor local switches in state. This incongruity could be solved by simply shifting the mechanism of state from the global signal to the local circuit. In this framework, traveling waves and neuromodulatory tone function as coordinating signals, instructing the state of distributed circuits. From this perspective, the fundamental unit of sleep and wake states might comprise non-overlapping libraries of spike patterns available to each circuit. A slow and global signal would generally coordinate circuits (Buzsáki and Schomburg, 2015; Varela et a., 2001; Buzsáki and Draguhn, 2004; Molle et al., 2006), thus determining the behavioral macrostate of the organism. Future work will be required to understand how fast embedding interacts with global signals such as neuromodulators.

Prior work in many species and brain regions describes shifts in neuronal activity that correlate with brain state (Noda and Adey, 1970; Abásolo et al., 2015; Watson et al., 2016; Levenstein et al., 2017; Brunwasser et al., 2022). However, such changes typically require extended observation, and while exhibiting statistically significant shifts across a large dataset, are too variable for use as a classification tool (Hengen et al., 2016; Xu et al., 2022). Here, small intervals of ensemble spiking activity are highly effective in separating each state (Fig. 6F). This is due to treating each neuron independently - it appears that, across many circuits, individual neurons are diverse in their state-dependent activity. While this is far more effective for classifying samples of sleep and wake than summary statistics, it requires a priori knowledge of each neuron’s profile, and the ability to track cells over extended periods of time. Despite this challenge, these data indicate that sleep and wake have complex effects on the activity of neighboring cells in the same microcircuit.

Based on behavior and low frequency rhythms, short states, namely microarousals and microsleeps, have received attention in both humans (Torsvall 1987; Halász 2005; Carskadon and Rechtschaffen, 2011) and rodents (Franken et al., 1998; Soltani et al., 2019). While there is disagreement regarding precise definitions (Kjaerby et al., 2022; Lüthi et al., 2022), these are common experiences, such as momentarily slipping into sleep when drowsy. Local states have also been described previously. Utilizing sleep deprivation, Vyazovskiy et al., (2011) demonstrated the ability of the waking cortex to support patches of slow wave activity, i.e. “local sleep”. This influential work led to the identification of wake-like and sleep-like oscillatory activity during sleep and wake, respectively, in humans (Nobili et al., 2011; Nir et al., 2011; Hung et al., 2013), as well as rats (Emrick et al., 2016). In our data, we found robust evidence of these previously described phenomena - each of which was based on low frequency rhythms. By excluding these events from our subsequent analyses of flickers, we demonstrate that the contribution of fast embedding to brain function is mechanistically distinct from brief and local alterations in low frequency rhythms. We were initially surprised to record all six possible types of flickers, given that some pairings of states do not map onto transitions observed under normal conditions, in particular, wake-to-REM flickers. However, similar transitions in the vertebrate brain are not impossible: in narcolepsy, for example, REM-like neural activity arises during wake (Kroeger & de Lecea, 2009; Cao & Guillemiault, 2017). Given the robust ability of CNNs to recover synthetic wake-to-REM flickers and their presence in all animals and all circuits, there is clearly a latent, high frequency signature of REM that can arise during waking. How this relates to the content of behavior remains unclear.

In this work, we take a neural-activity forward approach to discerning the minimal resolvable unit of brain state in anatomically and functionally diverse circuits. Two immediate approaches for future hold great promise. First, it will be fascinating to approach this problem in an unsupervised fashion; the number of reliably detectable substates may be far larger than appreciated. Second, a behavior-forward approach to state has great potential to further connect neuronal dynamics and brain function.

---

**Fig. S1.**
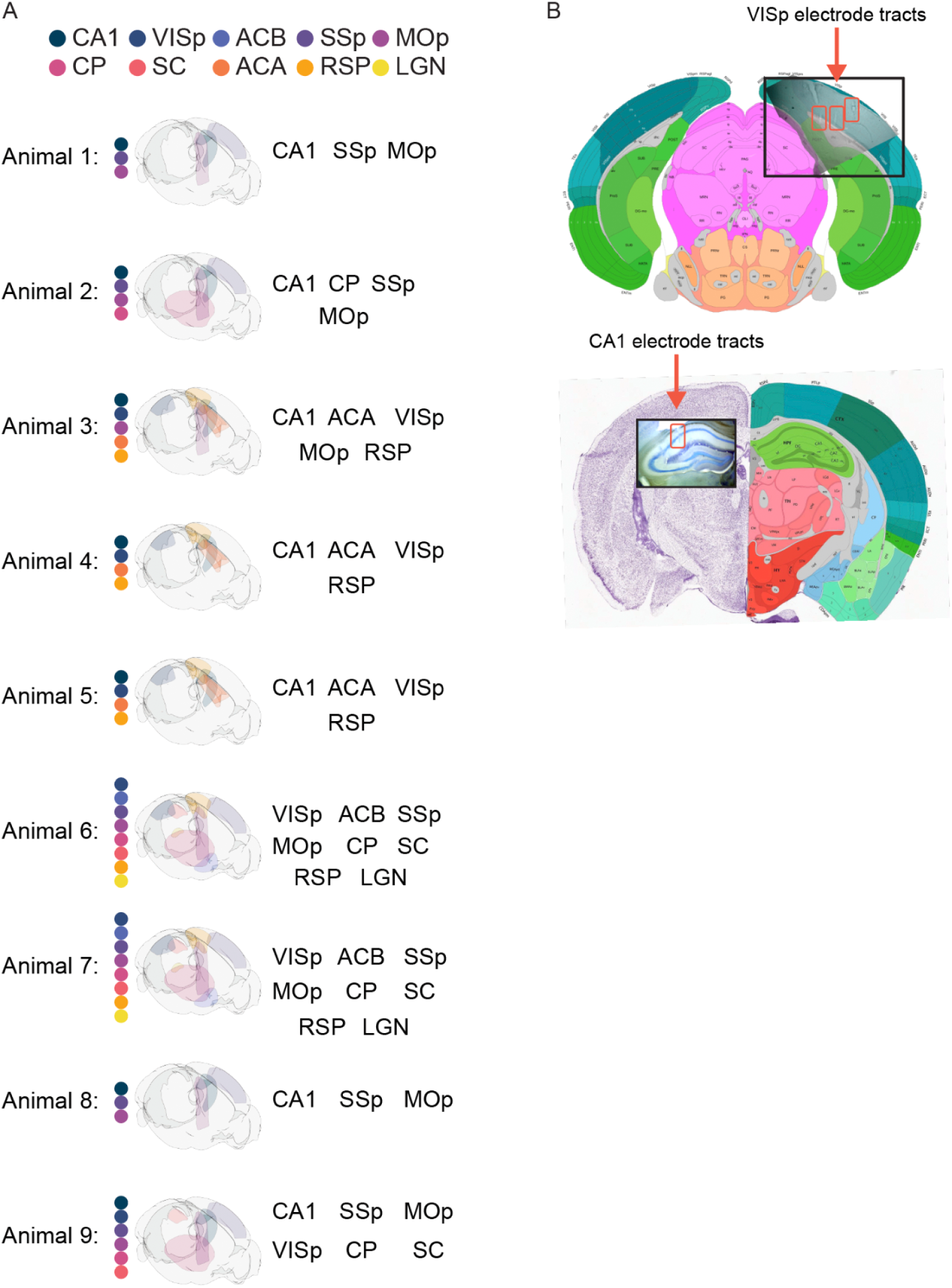
Individual animal implantation maps and electrode reconstruction. **A**, Representative illustration of the recording geometry in each animal used in this study. Each of nine mice were implanted with three to eight 64-channel tetrode bundles (total of 192 to 512 channels/animal). Colored dots correspond to the colored regions in each map. CA1: hippocampus, VISp: primary visual cortex, ACB: nucleus accumbens, SSp: primary somatosensory cortex, MOp: primary motor cortex, CP: caudoputamen, SC: superior colliculus, ACA: anterior cingulate, RSP: retrosplenial cortex, LGN: lateral geniculate nucleus thalamus (Claudi et al., 2020). **B,** Top- Example photomicrograph of tetrode tracts (red boxes) in primary visual cortex, aligned and overlaid on the corresponding coronal section from the Allen Mouse Brain Atlas (*Allen Institute for Brain Science*, 2012). Bottom- Example photomicrograph of CA1 hippocampus.

**Fig. S2.**
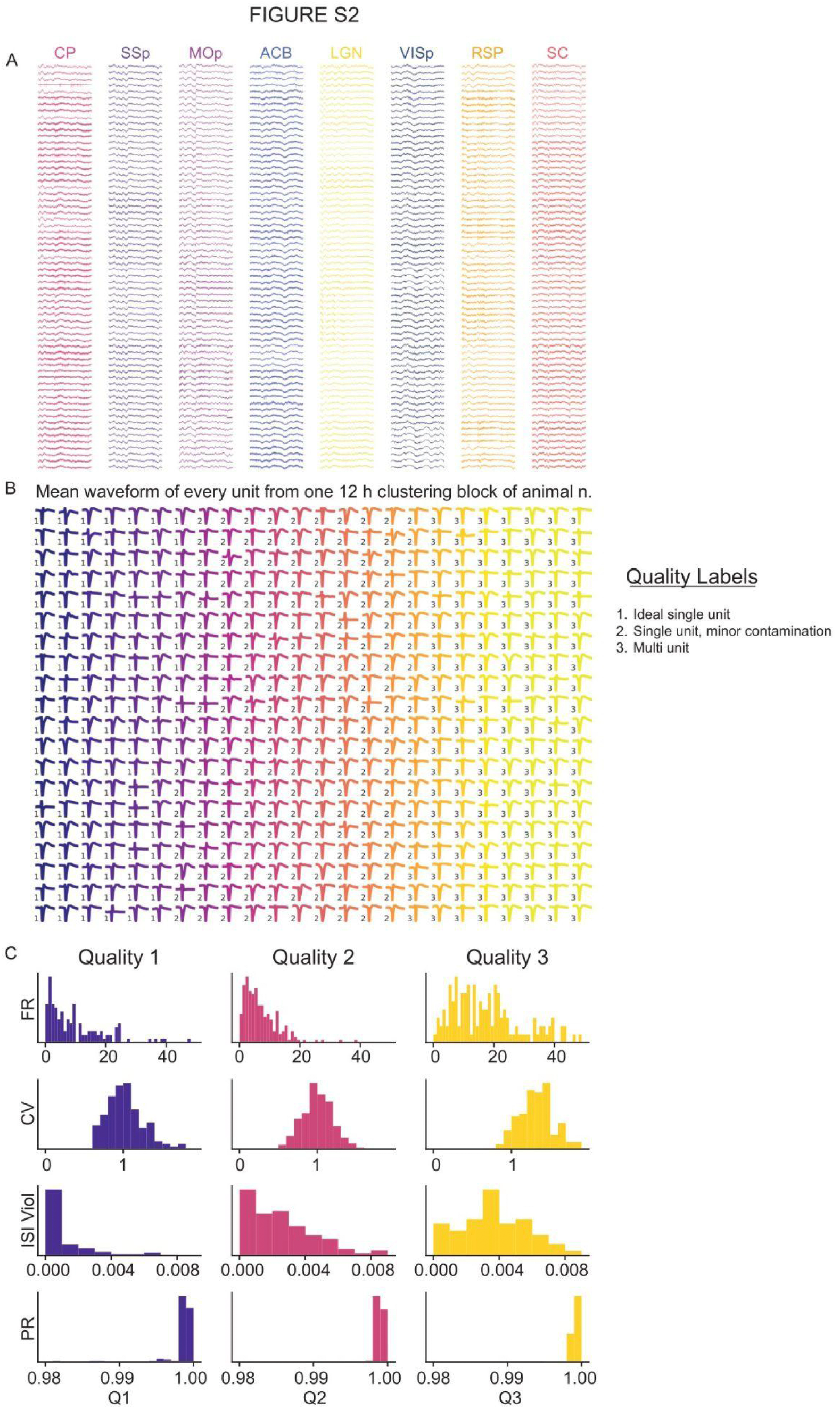
High-density multi-site recordings hold high-quality single-unit activity. Data shown is from Animal 9 roughly 10 weeks after implantation. **A,** This panel presents exemplary broadband traces from all 512 channels of recording. **B,** This illustrates the mean waveform from each spike-sorted unit. Number indicates the quality of each unit. Q1 and Q2 are single units. **C,** This panel showcases histograms of the distribution of various unit quality metrics across all units. In order: mean firing rate (FR), coefficient of variation (CV), portion of ISIs violating the minimum 1 ms refractory period (ISI Viol), presence ratio (PR).

**Fig. S3.**
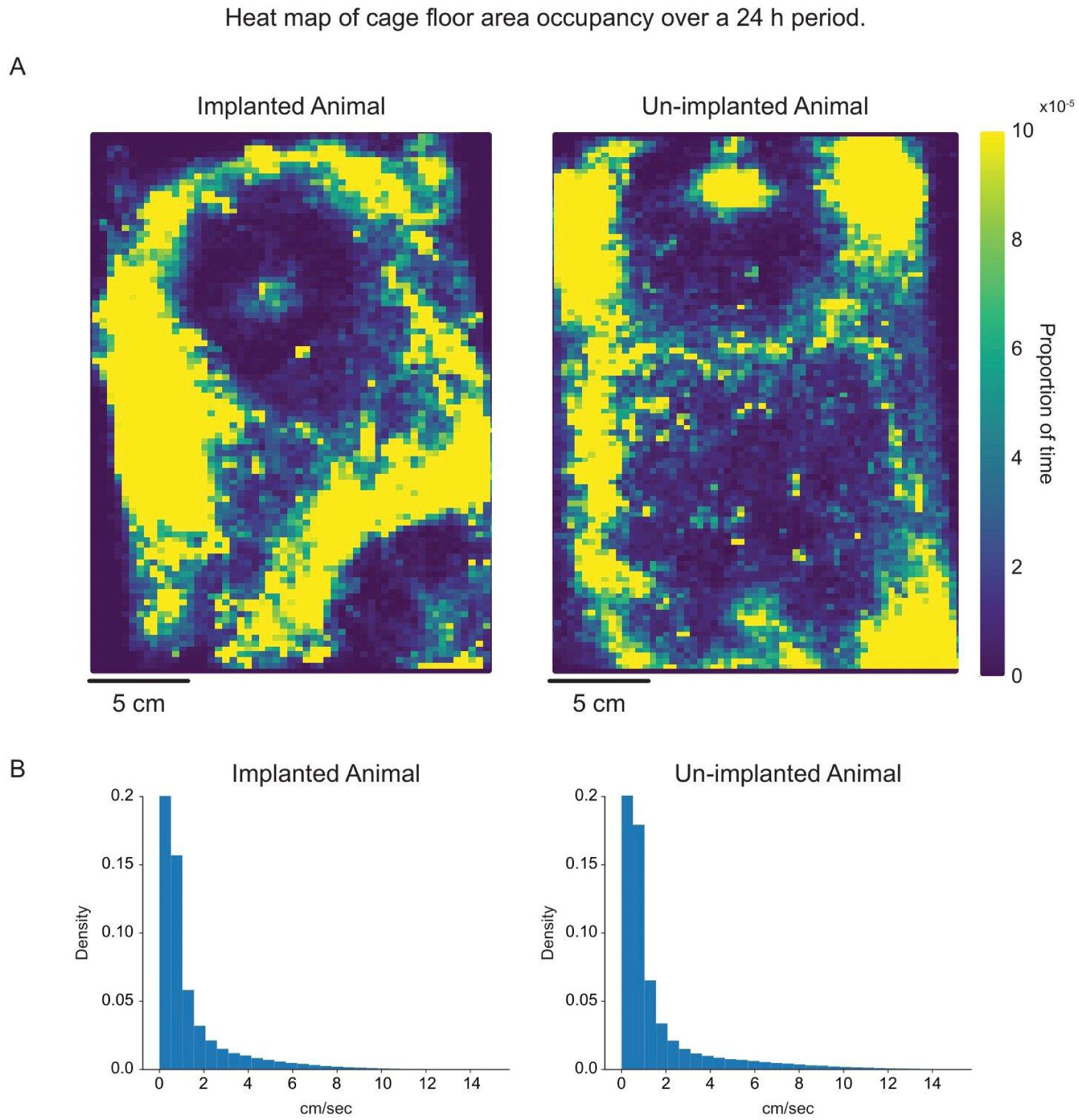
Implanted animals move freely. Implanted animal data is from Animal 3, and un-implanted animal data is from an age-matched control. **A,** Heatmap showing the portion of time spent in each portion of the cage floor by the animal over a 24 h period. Both animals explored 100% of the cage in this period. **B,** Histogram showing the distribution of velocities. Distributions are generally similar, but the unimplanted animal reached higher velocities while running. With a cutoff of 5 cm/s or better as locomotion, the mean locomotion velocity for the implanted mouse was 7.2 + /-2.3 cm/s (standard deviation) vs. the unimplanted mouse at 9.9 +/- 4.0cm/s. The difference was apparent largely in the upper bounds of locomotion speed. The 95th percentile of speed for the implanted mouse was 11.4 cm/s vs. 17.6 cm/s for the unimplanted mouse.

**Fig. S4.**
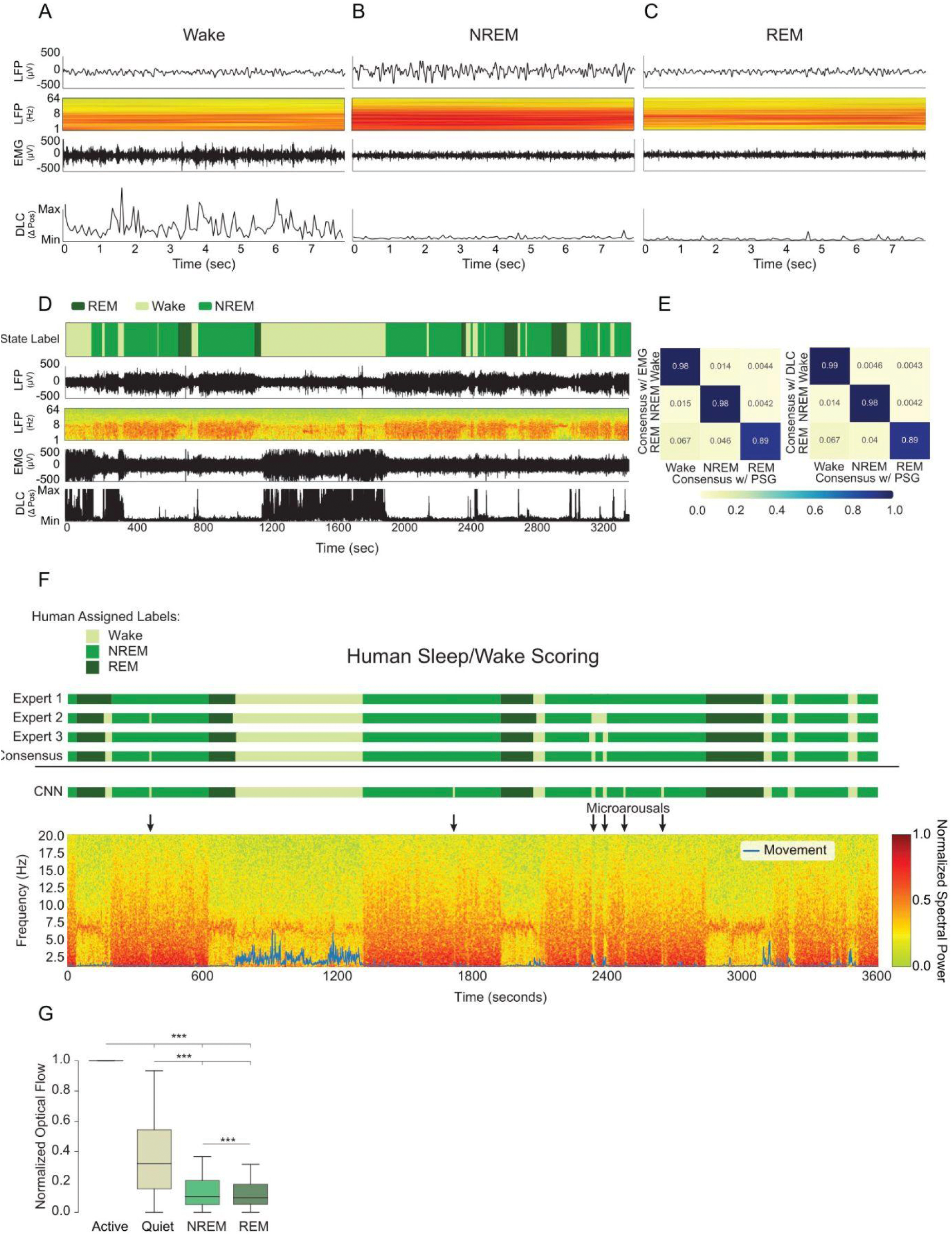
Human sleep scoring, polysomnography, example traces. **A**, 8-seconds of exemplary polysomnography data from an animal during wake. From top-to-bottom: the low-passed LFP trace, the LFP spectrogram, the EMG, and the MinMax normalized change in position tracked by DeepLabCut (DLC) are shown (Mathis et al., 2018). To form EEG-like traces, raw data were low-pass filtered at 125 Hz, downsampled to 500 Hz, and 8 non-adjacent channels were averaged. **B,** 8-seconds of exemplary data from NREM, **C,** 8 seconds of exemplary data from REM. **D,** An hour of data sleep-scored for Wake, NREM and REM with polysomnography. Delta band power (0.1-4 Hz) is highly enriched in NREM (slow wave) sleep, theta band power (6-8 Hz) is enriched in REM sleep, and high resolution motor output disambiguates periods of waking, including microarousals (Watson et al., 2016). **E,** Confusion matrices for average individual scorer performance using EMG or DLC for motion (rows) to consensus scoring among three expert scorers using polysomnography, including DLC and EMG (columns). **F,** Exemplary hour of data sleep-scored by three experts to produce a consensus score, which is used as training data for a CNN (predictions shown). Blue movement trace is based on DeepLabCut. **G,** Box plots of normalized optical flow (see methods section “Flickers During Waking Behavior”) for active wake (active motion state), quiet wake (inactive motion state), NREM, and REM. Highly significant differences in means were observed for all comparisons (p<0.001, ANOVA with post-hoc EMMeans with Tukey correction).

**Fig. S5.**
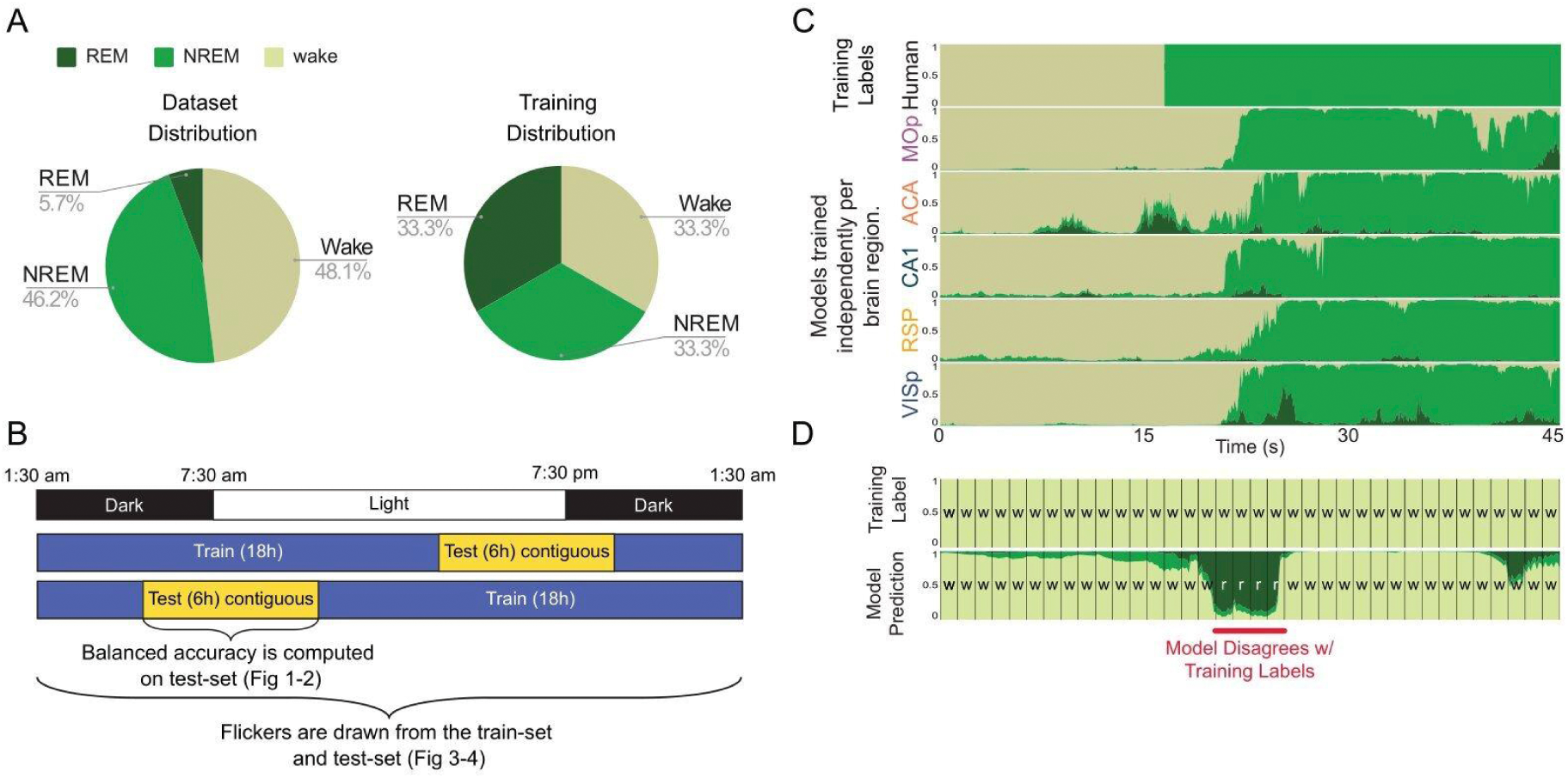
CNN train/test separation, model independence, and flickering. **A**, The three arousal states, REM, NREM, and wake, were not evenly distributed in the recorded data (dataset distribution). Training on equal amounts of data from each state, i.e., class balancing (training distribution), prevents CNN models from simply learning to predict the most frequent class whenstate information is not clear. **B,** 24 h of data spanning a complete 12-12 light/dark cycles was included from each animal. Cartoon is a schematic illustrating 24 h of light/dark (top row), and two examples of train/test segmentation of data (bottom rows). CNNs were trained on 18 h of data. Test data comprised six contiguous hours of data spanning a light/dark transition. **C,** Models were trained using consensus human labels (top row). Independent models were trained and tested in each brain region within each animal (bottom five rows show the predictions of five models, each trained to recapitulate the human labels from data recorded within the brain region indicated on the left). CNNs are well suited to overcome label error, such as when a human score lags or leads a transition. The provided example demonstrates disagreement and model independence surrounding a global state transition (wake into NREM). The y-axis of CNN models indicates instantaneous confidence [0-1] in each state via the proportion of each of three colors at each point in time. **D,** In contrast to assessment of model accuracy, flickers are extracted from both train and test output. Flickers detected in the training component of data represent CNNs directly disagreeing with training labels (red line indicates example of wake-to-REM flicker). Human labeled data (top row) and corresponding CNN predictions (bottom row) are computed on an interval set by the experimenter. Flickers were extracted and cross validated in both 2.6 s and 1s interval models.

**Fig. S6.**
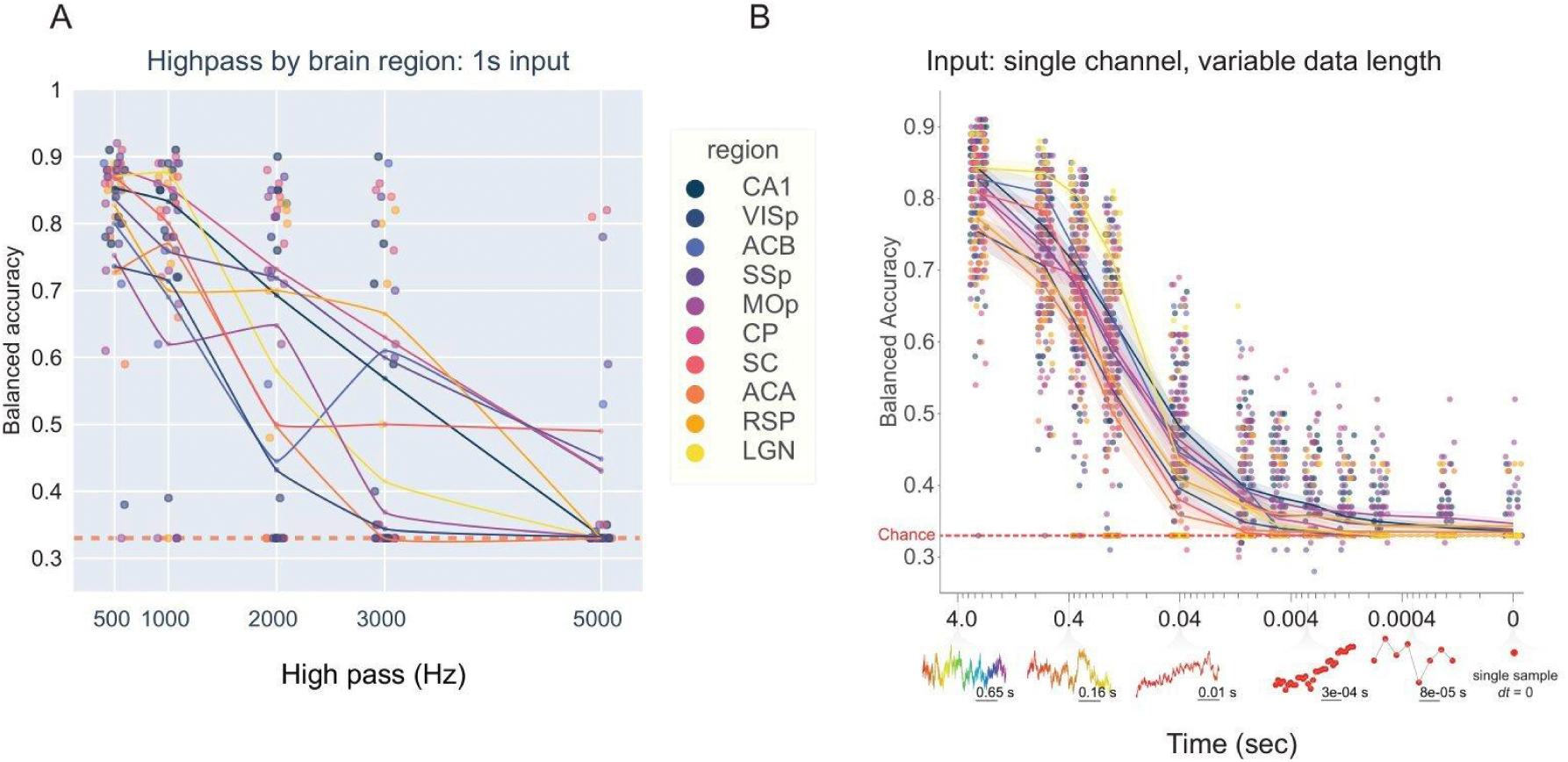
CNN accuracy by region as a function of high-pass and sample size reduction. **A**, Balanced accuracy of CNNs trained and tested on progressively high-passed raw data from each recorded brain region (n=45 implants, 9 animals, 10 regions). Dot color represents the region from which this model was trained. The region-colored line traces the average balanced accuracy of models trained on data from that region across various levels of high-pass. High-pass filtering significantly decreased brain state information above 1,000 Hz (y = -9.892e-05*x+ 0.84, p<0.001, r2 = 0.43). **B,** To directly test the minimum time interval in which sleep and wake states reliably structure neural dynamics, we trained and tested a series of CNNs on single channel data, each model operating on a progressively smaller interval of data (from 2.6 s to 0 s). Accuracy declined as a function of number of input sample points (pearson correlation: r=0.650464, p<0.001). Example data at various input sizes is shown below the x-axis. Model accuracy is shown as a function of region (marker color). Regional means are shown in colored lines (+/- SEM, shaded area). To examine inter-regional differences at these small time scales, we performed a linear mixed effects regression where balanced accuracy is a function of brain region and sample size (implant included as a random effect).

**Fig. S7.**
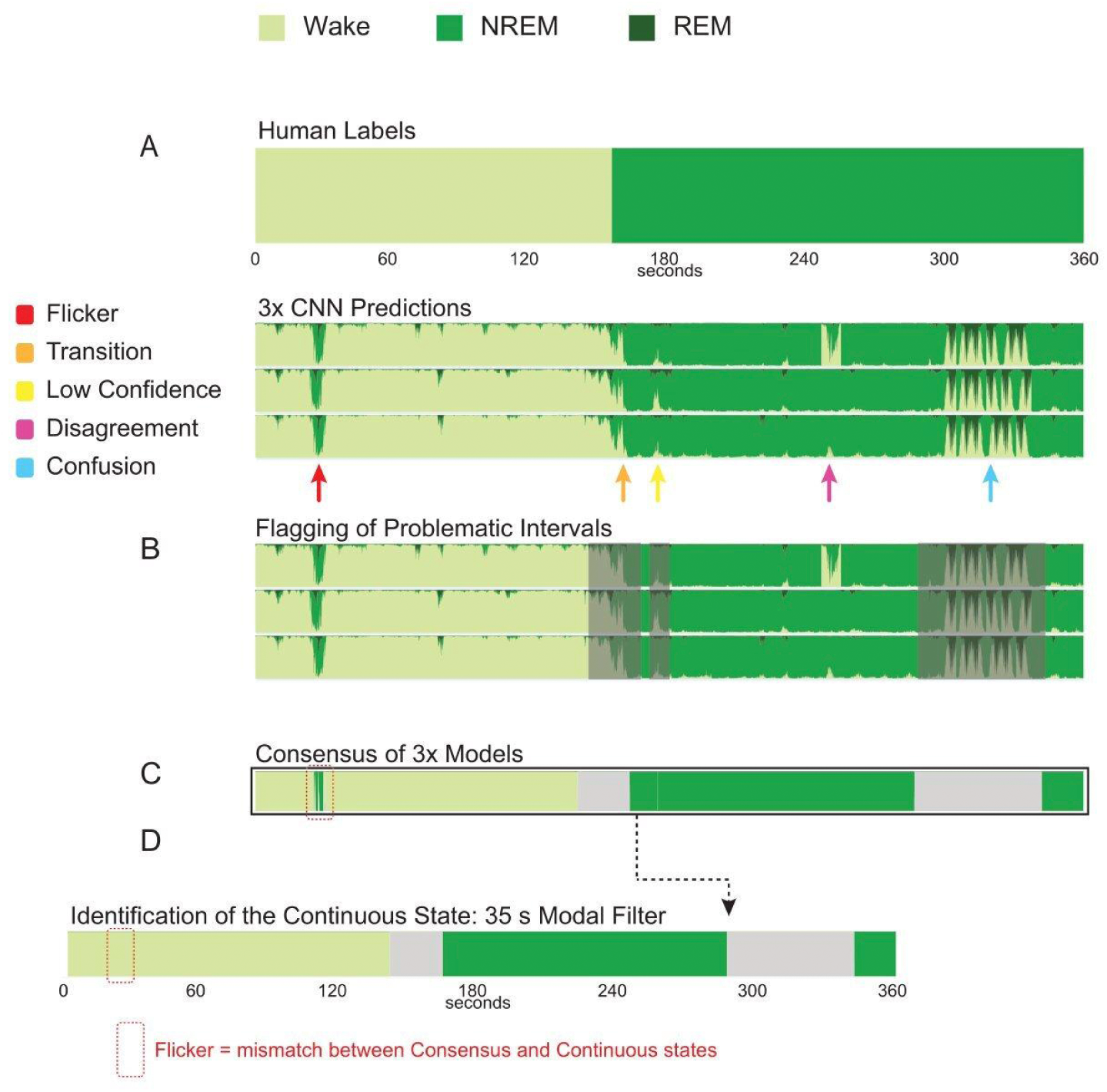
Flicker definition method applied to synthetic data. To illustrate the process of identifying flickers within data, we generated a synthetic dataset equivalent to 64 channels of recording from a single brain region. The synthetic data were designed to contain examples of all forms of noise that our algorithms exclude in the process of identifying flickers. **A,** Human experts score animal arousal state based on polysomnography (top). Based on these scores, three CNNs are trained to identify NREM, REM and wake based on raw neural data (1 s input) from all 64 channels within a brain region. Shown are triplicate CNN-generated state scores for our simulated brain region. **B,** We identify and exclude two forms of extended intervals during which flickers are not to be considered. First, we identify windows of time during which a *transition* (orange arrow) between two sustained states occurs. Second, we identify rare instances of label *confusion* (typically the result of extreme noise or CNN error) (blue arrow). This is achieved by automatically excluding any 35 s epochs with a mean confidence <75%. To exclude low confidence *noise* (yellow arrow) from our analyses, we eliminate 1s epochs with a mean confidence <75%. For each time point, we then assign the label of the most confidently-predicted state. **C,** To exclude *disagreements* (magenta arrow), or artifacts of particular CNN models (i.e. predicted by only one of three CNNs), we collapse predictions across models by selecting the most commonly predicted state at each time point. **D,** We then slide a 35 s modal filter across the majority state array to create a label corresponding to the stable macro state surrounding each point in time. *Flickers* are defined as disagreements between the consensus array and the continuous array (red arrow).

**Fig. S8.**
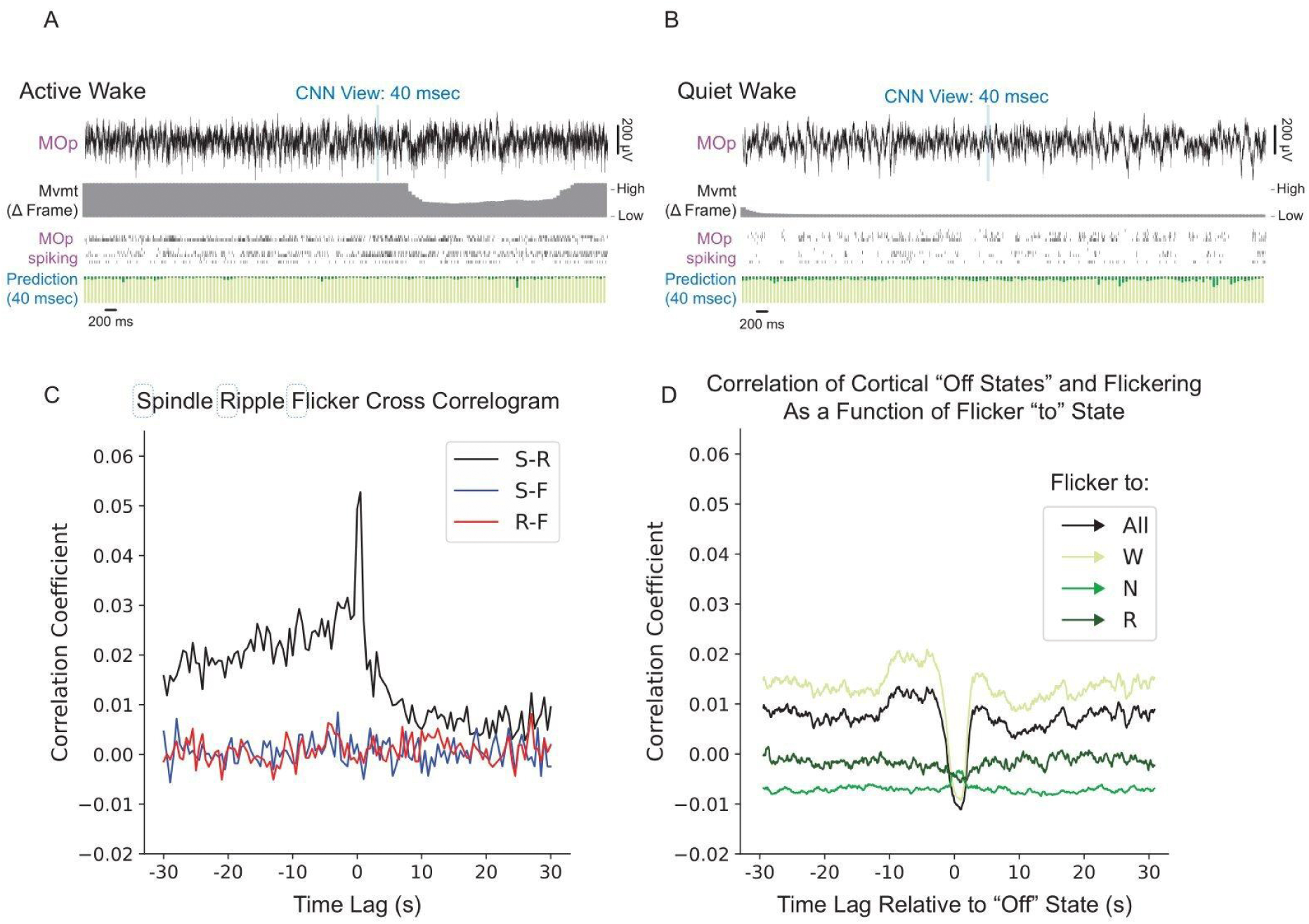
40 ms CNNs are resilient to oscillatory substates. **A**, Top- Broadband trace of exemplary activity during active wake in MOp over several seconds. Blue box shows the width of an individual input data used by the 40 ms CNN to predict state. Middle- Raster of MOp spiking. Bottom- Stacked barplot of CNN prediction probabilities across the three states every 1/15 s. **B,** Exemplary data from MOp during quiet wake. **C,** Cross-correlogram between spindles (S), ripples (R), and flickers (F) in Animal 5. A strong central peak in the cross-correlogram is observed between spindles and ripples consistent with prior work (Siapas & Wilson, 1998). No significant positive correlation is observed with flickers by spindles or ripples. **D)** Cross-correlogram between OFF-states and flickers of various states (for instance ->W includes NREM-to-wake, and REM-to-wake). A significant central correlation trough is observed in flickers to wake, meaning flickering is reduced during sleep OFF states.

**Fig. S9.**
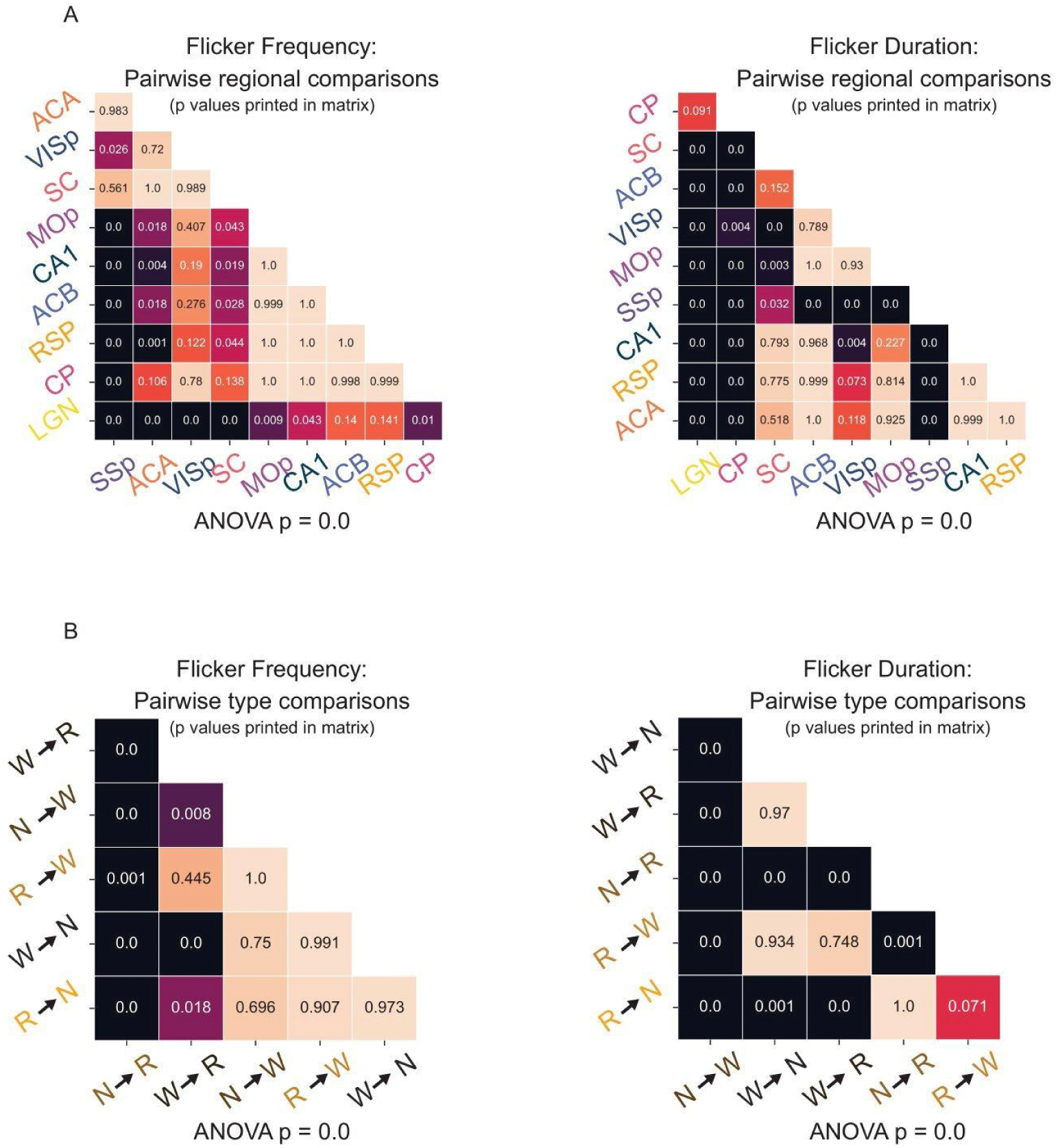
Flicker timing shows significant differences by circuit and flicker type. Pairwise comparisons between subsets of flickers using EMMeans with Tukey correction as a post hoc-test for an ANOVA fit to a linear mixed effects model with animal as a random effect. For all ANOVAs, p<0.001. p-Values for the pairwise comparison of row and column are printed inside a cell of the diagonal matrices. **A,** Left- Significant differences in flicker frequency (i.e. rate) when grouped by region (i.e. circuit). Right- Significant differences in flicker duration when grouped by region. **B,** Left- Significant differences in flicker frequency (i.e. rate) when grouped by flicker type. Right- Significant differences in flicker duration when grouped by flicker type.

**Fig. S10.**
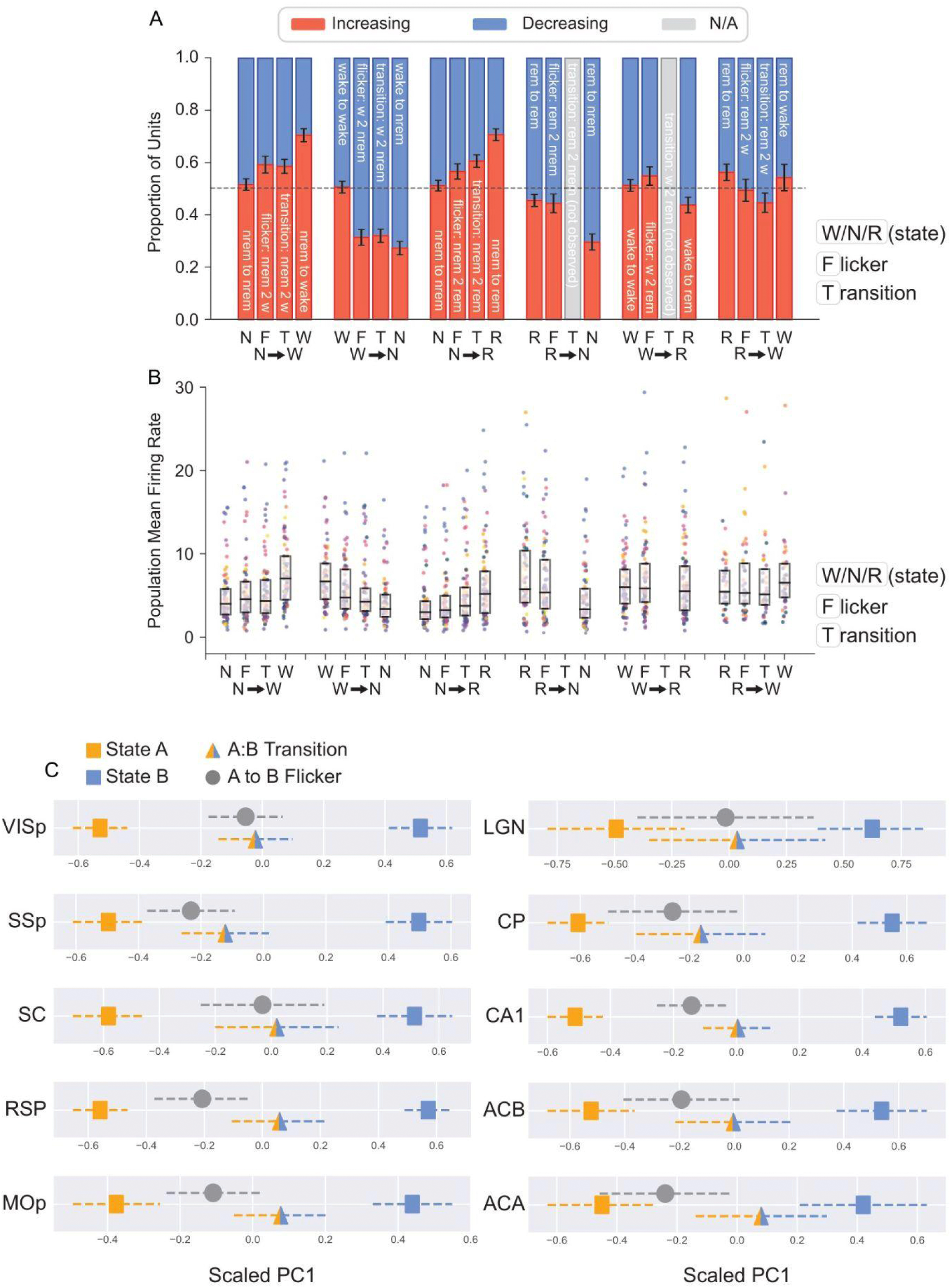
Flickering corresponds to transition-like spiking patterns. This figure expands on trends in spiking presented in Fig. 6. **A,** The portion of units whose sampled instantaneous firing rate was different relative to a random sample of the surrounding state: surrounding state vs. surrounding state (negative control), surrounding state-to-predicted state flicker, surrounding state-to-predicted state transition, and surrounding state vs. predicted state. **B,** The mean single unit firing rate of recorded circuits during a surrounding state, surrounding state-to-predicted state flicker, surrounding state-to-predicted state transition, and predicted state. Box plots show the intermediate quartiles with outliers as swarm scatter colored by region. **F)** Mean scaled PC1 projections for the ten regions. For each region surrounding state, predicted flicker state, flicker, and transition are shown. Error bars are the SEM multiplied by square-root of degrees of freedom (number of animals). See Supplemental Tables 2-5 for significance of pairwise comparisons based on linear mixed effects model projection ∼ sample type (i.e. flicker, transition, surrounding, predicted) + (1 | animal) with post-hoc EMMeans with Tukey correction. Bonferroni multiple hypothesis correction was applied based on the number of regions.

**Table S1.** Number of spike-sorted units. The number of single-units (included) and multi-units spike-sorted from a one-hour block for each recorded circuit in each animal.

| Animal | Region | Single-Units | Multi-Units |
| --- | --- | --- | --- |
| 1 | MOp | 11 | 21 |
| 1 | CA1 | 31 | 14 |
| 1 | SSp | 19 | 9 |
| 2 | SSp | 13 | 23 |
| 2 | MOp | 13 | 31 |
| 2 | CA1 | 16 | 30 |
| 2 | ACB | 15 | 30 |
| 3 | MOp | 10 | 21 |
| 3 | ACA | 33 | 40 |
| 3 | CA1 | 12 | 27 |
| 3 | RSP | 39 | 28 |
| 3 | VISp | 16 | 23 |
| 4 | ACA | 14 | 18 |
| 4 | RSP | 45 | 24 |
| 4 | VISp | 36 | 17 |
| 4 | CA1 | 30 | 15 |
| 5 | ACA | 8 | 4 |
| 5 | RSP | 18 | 16 |
| 5 | VISp | 8 | 4 |
| 6 | CP | 18 | 18 |
| 6 | SSp | 41 | 8 |
| 6 | MOp | 28 | 12 |
| 6 | ACB | 29 | 18 |
| 6 | LGN | 7 | 4 |
| 6 | VISp | 25 | 13 |
| 6 | RSP | 40 | 23 |
| 6 | SC | 15 | 7 |
| 7 | MOp | 42 | 12 |
| 7 | SSp | 21 | 22 |
| 7 | CP | 18 | 27 |
| 7 | ACB | 41 | 20 |
| 7 | LGN | 63 | 19 |
| 7 | SC | 8 | 12 |
| 7 | VISp | 14 | 11 |
| 8 | SSp |  | 5 |
| 8 | CA1 | 16 | 11 |
| 8 | MOp | 39 | 20 |
| 8 | MOp | 19 | 17 |
| 9 | CP | 38 | 29 |
| 9 | CP | 35 | 16 |
| 9 | MOp | 20 | 16 |
| 9 | CA1 | 30 | 14 |
| 9 | CA1 | 16 | 20 |
| 9 | SSp | 29 | 14 |
| 9 | SC | 17 | 25 |

**Table S2.** ANOVA results for PCA analysis of flicker types. ANOVA of linear mixed effects model projection ∼ sample type (i.e. transition, flicker, etc.) + (1 | animal). Bonferroni multiple hypothesis correction was applied based on the number of flicker types.

| Flicker Type | F value | Pr(>F) |
| --- | --- | --- |
| wake - to - NREM | 134.276099 | 7.43E-53 |
| wake - to – REM | 299.771071 | 3.81E-64 |
| NREM – to – wake | 154.089017 | 5.77E-59 |
| NREM – to – REM | 190.147826 | 1.78E-69 |
| REM – to – wake | 73.267739 | 7.91E-28 |
| REM – to – NREM | 401.32479 | 1.95E-68 |

**Table S3.**
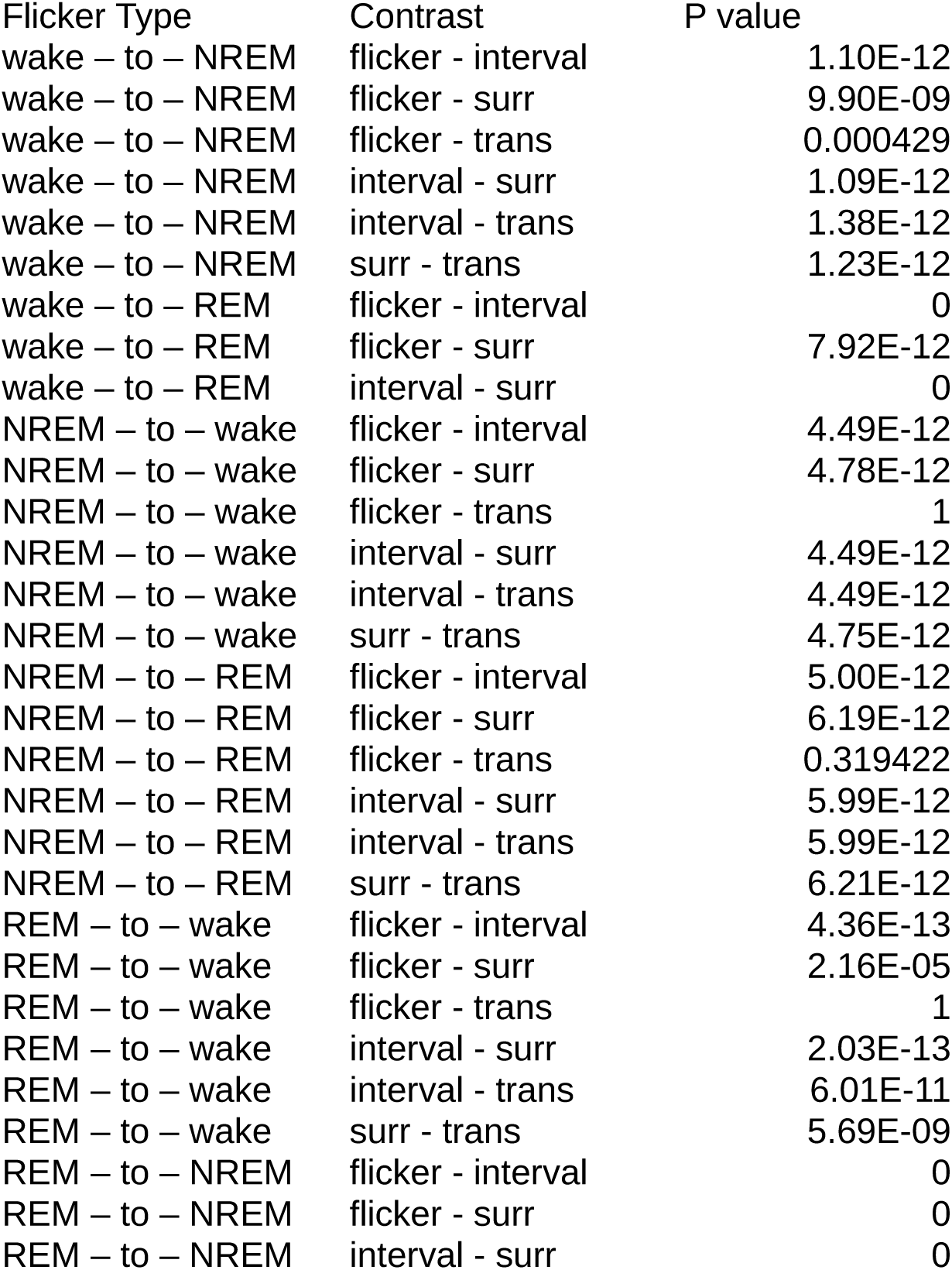
Pairwise comparisons for PCA analysis of flicker types. ANOVA of linear mixed effects model projection ∼ sample type (i.e. transition, flicker, etc.) + (1 | animal). Result of post-hoc EMMeans with Tukey correction. Bonferroni multiple hypothesis correction was applied based on the number of flicker types.

| Flicker Type | Contrast | P value |
| --- | --- | --- |
| wake – to – NREM | flicker - interval | 1.10E-12 |
| wake – to – NREM | flicker - surr | 9.90E-09 |
| wake – to – NREM | flicker - trans | 0.000429 |
| wake – to – NREM | interval - surr | 1.09E-12 |
| wake – to – NREM | interval - trans | 1.38E-12 |
| wake – to – NREM | surr - trans | 1.23E-12 |
| wake – to – REM | flicker - interval | 0 |
| wake – to – REM | flicker - surr | 7.92E-12 |
| wake – to – REM | interval - surr | 0 |
| NREM – to – wake | flicker - interval | 4.49E-12 |
| NREM – to – wake | flicker - surr | 4.78E-12 |
| NREM – to – wake | flicker - trans | 1 |
| NREM – to – wake | interval - surr | 4.49E-12 |
| NREM – to – wake | interval - trans | 4.49E-12 |
| NREM – to – wake | surr - trans | 4.75E-12 |
| NREM – to – REM | flicker - interval | 5.00E-12 |
| NREM – to – REM | flicker - surr | 6.19E-12 |
| NREM – to – REM | flicker - trans | 0.319422 |
| NREM – to – REM | interval - surr | 5.99E-12 |
| NREM – to – REM | interval - trans | 5.99E-12 |
| NREM – to – REM | surr - trans | 6.21E-12 |
| REM – to – wake | flicker - interval | 4.36E-13 |
| REM – to – wake | flicker - surr | 2.16E-05 |
| REM – to – wake | flicker - trans | 1 |
| REM – to – wake | interval - surr | 2.03E-13 |
| REM – to – wake | interval - trans | 6.01E-11 |
| REM – to – wake | surr - trans | 5.69E-09 |
| REM – to – NREM | flicker - interval | 0 |
| REM – to – NREM | flicker - surr | 0 |
| REM – to – NREM | interval - surr | 0 |

**Table S4.** ANOVA results for PCA analysis of regions. ANOVA of linear mixed effects model projection ∼ sample type (i.e. transition, flicker, etc.). Bonferroni multiple hypothesis correction was applied based on the number of regions.

| Region | F value | Pr(>F) |
| --- | --- | --- |
| CA1 | 170.920265 | 4.37E-57 |
| VISp | 139.313237 | 2.09E-45 |
| SSp | 102.302738 | 2.23E-39 |
| MOp | 68.711296 | 2.66E-31 |
| CP | 82.351533 | 5.13E-25 |
| SC | 68.113438 | 2.92E-23 |
| ACA | 39.807623 | 1.12E-16 |
| RSP | 159.480878 | 9.83E-45 |
| LGN | 28.355577 | 3.80E-10 |
| ACB | 59.913145 | 1.73E-20 |

**Table S5.**
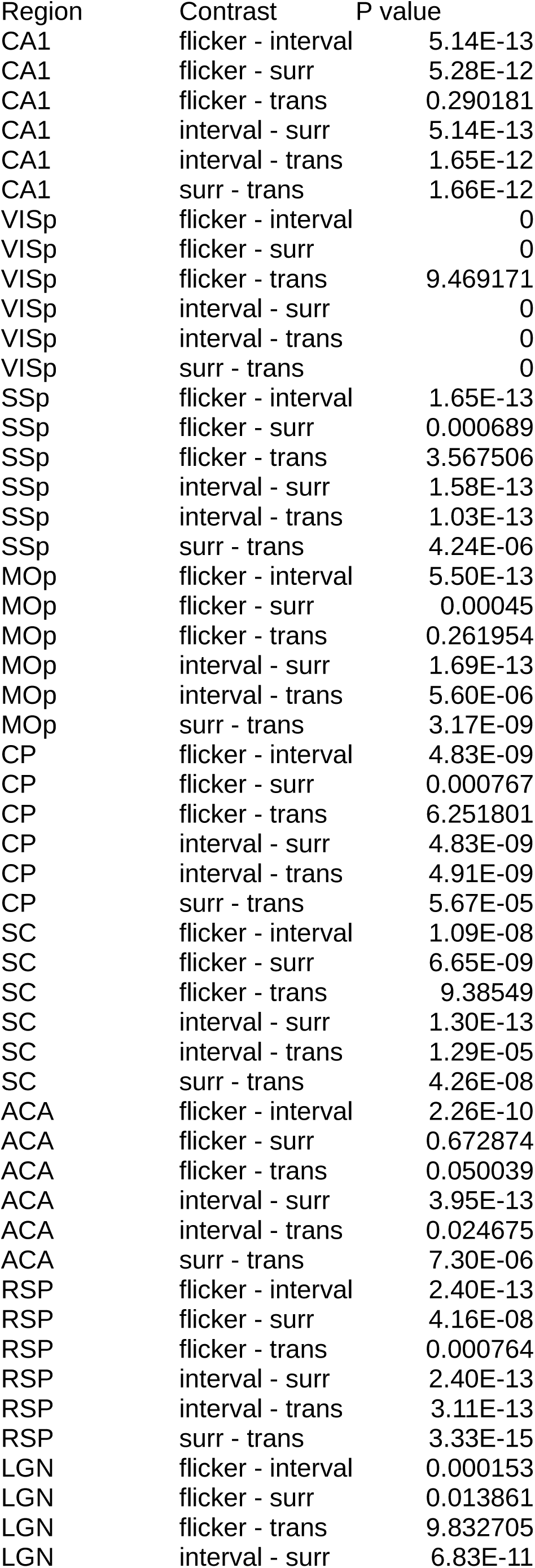
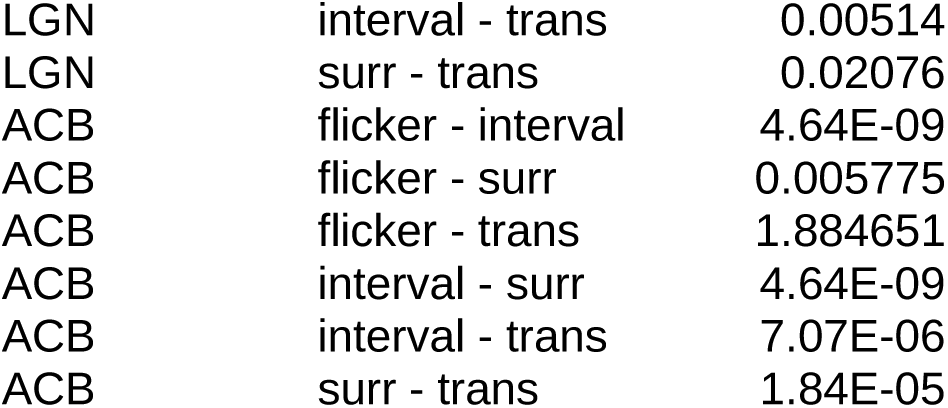
Pairwise comparisons for PCA analysis of regions. ANOVA of linear mixed effects model projection ∼ sample type (i.e. transition, flicker, etc.) + (1 | animal). Result of post-hoc EMMeans with Tukey correction. Bonferroni multiple hypothesis correction was applied based on the number of regions.

| Region | Contrast | P value |
| --- | --- | --- |
| CA1 | flicker - interval | 5.14E-13 |
| CA1 | flicker - surr | 5.28E-12 |
| CA1 | flicker - trans | 0.290181 |
| CA1 | interval - surr | 5.14E-13 |
| CA1 | interval - trans | 1.65E-12 |
| CA1 | surr - trans | 1.66E-12 |
| VISp | flicker - interval | 0 |
| VISp | flicker - surr | 0 |
| VISp | flicker - trans | 9.469171 |
| VISp | interval - surr | 0 |
| VISp | interval - trans | 0 |
| VISp | surr - trans | 0 |
| SSp | flicker - interval | 1.65E-13 |
| SSp | flicker - surr | 0.000689 |
| SSp | flicker - trans | 3.567506 |
| SSp | interval - surr | 1.58E-13 |
| SSp | interval - trans | 1.03E-13 |
| SSp | surr - trans | 4.24E-06 |
| MOp | flicker - interval | 5.50E-13 |
| MOp | flicker - surr | 0.00045 |
| MOp | flicker - trans | 0.261954 |
| MOp | interval - surr | 1.69E-13 |
| MOp | interval - trans | 5.60E-06 |
| MOp | surr - trans | 3.17E-09 |
| CP | flicker - interval | 4.83E-09 |
| CP | flicker - surr | 0.000767 |
| CP | flicker - trans | 6.251801 |
| CP | interval - surr | 4.83E-09 |
| CP | interval - trans | 4.91E-09 |
| CP | surr - trans | 5.67E-05 |
| SC | flicker - interval | 1.09E-08 |
| SC | flicker - surr | 6.65E-09 |
| SC | flicker - trans | 9.38549 |
| SC | interval - surr | 1.30E-13 |
| SC | interval - trans | 1.29E-05 |
| SC | surr - trans | 4.26E-08 |
| ACA | flicker - interval | 2.26E-10 |
| ACA | flicker - surr | 0.672874 |
| ACA | flicker - trans | 0.050039 |
| ACA | interval - surr | 3.95E-13 |
| ACA | interval - trans | 0.024675 |
| ACA | surr - trans | 7.30E-06 |
| RSP | flicker - interval | 2.40E-13 |
| RSP | flicker - surr | 4.16E-08 |
| RSP | flicker - trans | 0.000764 |
| RSP | interval - surr | 2.40E-13 |
| RSP | interval - trans | 3.11E-13 |
| RSP | surr - trans | 3.33E-15 |
| LGN | flicker - interval | 0.000153 |
| LGN | flicker - surr | 0.013861 |
| LGN | flicker - trans | 9.832705 |
| LGN | interval - surr | 6.83E-11 |
| LGN | interval - trans | 0.00514 |
| LGN | surr - trans | 0.02076 |
| ACB | flicker - interval | 4.64E-09 |
| ACB | flicker - surr | 0.005775 |
| ACB | flicker - trans | 1.884651 |
| ACB | interval - surr | 4.64E-09 |
| ACB | interval - trans | 7.07E-06 |
| ACB | surr - trans | 1.84E-05 |

## Methods

### Mice

All procedures involving mice were performed in accordance with protocols approved by the Washington University in Saint Louis Institutional Animal Care and Use Committee, following guidelines described in the US National Institutes of Health Guide for the Care and Use of Laboratory Animals. C57BL/6 mice from Charles River were used. Seven female and two male mice were used in this study. Mice were at least 100 d old at the beginning of recording (mean age 220 d). Mice were housed in an enriched environment and kept on a 12:12 h light:dark cycle. Mice had *ad libitum* access to food and water.

### Surgery

All mice underwent multisite electrode array implantation surgery. Mice were anesthetized with isoflurane (1-2% in air) and administered slow release buprenorphine (ZooPharm, 0.1 mg kg^-1^). The mouse’s skull was secured in a robotic stereotaxic instrument (NeuroStar, Tubingen, Germany), and the skin and periosteum covering the dorsal surface of the skull was removed. Pitch, yaw, and roll were calculated to maximize alignment with stereotaxic atlases. Three to eight craniotomies (diameter 1-1.5 mm) were drilled using the automatic drilling function of the stereotaxic robot, and the dura was resected. In each animal, each of three to eight brain regions were implanted with a custom 64-channel tetrode-based array. Arrays were fixed (not drivable) and separated from headstage hardware by a flex cable, thus allowing an arbitrary geometry of multiple probes. Across nine mice, a total of 45 implants spanned 10 unique brain regions. Coordinates were as follows (AP/ML/DV relative to bregma and dura, in mm): CA1 (n = 6, -2.54/-1.75/-1.5), VISp (n = 5, -3.8/-2.8/-0.8), ACB (n = 2, 1.25/-0.81/-3.94), SSp (n = 6, -0.5/2.25/-0.8), MOp (n = 7, 1.25/1.75/-1), CP (n = 4, 0.5/1.52/-2.56), SC (n = 3, -4.0/-1.0/-1.0), ACA (n = 3, 0.6/0.8/-0.94), RSP (n = 4, -1.5/0.3/-0.9), and LGN (n = 2, -2.25/2.26/-2.40).

Electrode bundles were lowered into brain tissue at a rate of 5mm/minute using a custom built stereotaxic vacuum holder. Anatomical location was confirmed post hoc via histological reconstruction (Extended Data Fig.1b). Arrays were secured with dental cement (C&B-Metabond Quick! Luting Cement, Parkell Products Inc; Flow-It ALC Flowable Dental Composite, Pentron), and headstage electronics (eCube, White Matter LLC) were bundled and secured in custom 3D-printed housing. Eight-module (512 channel) implants including arrays, cement, and headstage materials weighed approximately 4g. Mice were administered meloxicam (Pivetal, 5 mg kg^-1^ day^-1^ for three days) and dexamethasone (0.5 mg kg^-1^ day^-1^ for three days) and allowed to recover in the recording chamber for at least one week prior to recording.

### Recording

Recordings were made using custom tetrode-based arrays. 16 tetrodes (64 channels) were soldered to a custom-designed PCB (5 mm x 5 mm x 200 μm) which stacked horizontally with a similarly sized amplifier chip. PCB/amplifier pairs further stacked with up to 7 additional pairs (total 8 modules, 512 channels). Recordings were conducted in an enriched home cage environment with social access to a litter mate through a perforated acrylic divider. Freely behaving mice were attached to a custom built cable with in-line commutation. Neuronal signals were amplified, digitized, and sampled at 25 kHz alongside synchronized 15-30 fps video using the eCube Server electrophysiology system (White Matter LLC). Recordings were conducted continuously for between two weeks and three months. Data and video were continuously monitored using Open Ephys (Siegle et al., 2017) and Watch Tower (White Matter LLC). 24 h blocks of data were identified for inclusion in these studies first by the absence of hardware and/or software problems, for example cable disconnects or dropped video frames, respectively, and second by the maximal yield of active channels. Beyond these criteria, selection of 24 h blocks was arbitrary. The same 24 h block was utilized for all recorded circuits in an individual animal.

For experiments involving spike-sorted data, raw data were bandpass filtered between 350 and 7,500 Hz and spike waveforms were extracted and clustered using a modified version of SpikeInterface (Buccino et al., 2020) and MountainSort4 (Chung et al., 2017) with curation turned off. A custom XGBoost was used to identify those clusters constituting single units. Clusters identified as single units were manually inspected to confirm the presence of high amplitude spiking, stable spike amplitude over time, consistent waveform shapes, and little to no refractory period contamination.

### Probe Localization

Following recording, mice were perfused with 4% formaldehyde (PFA), the brain was extracted and immersion fixed for 24h at 4° C in PFA. Brains were then transferred to a 30% sucrose solution in PBS and stored at 4° C until brains sank. Brains were then sectioned at 50 μm on a cryostat. Sections were rinsed in PBS prior to mounting on charged slides (SuperFrost Plus, Fisher) and stained with cresyl violet. Stained sections were aligned with the Allen Institute Mouse Brain Atlas (*Allen Institute for Brain Science*, 2012) and tetrode tracks were identified under a microscope (Extended Data Fig. 1B).

### Consensus Sleep Scoring

Three experts independently scored the arousal state of each mouse using custom software (Extended Data Fig. 4D). Briefly, the LFP spectral power (0.1-60 Hz) was extracted from five channels selected from cortical implantations and averaged. Movement data was extracted from video recordings using DeepLabCut (Mathis et al., 2018) or EMG. LFP and movement data were preliminarily sleep-scored in 4 s epochs by a random forest. Human experts then evaluated LFP spectral density, movement, and random forest output in 4 s epochs. Scoring software provided immediate access to temporally aligned video for disambiguation.

The three independently generated arrays of state assignments were then compared and all disagreements were identified. The three contributing human experts then, as a group, reevaluated each epoch of disagreement and generated a consensus state label.

### Convolutional Neural Network Construction and Experiments

In this work, we used a single neural network model. We chose this architecture as it is particularly robust to label error, and is thus well suited to tasks in which there may be substantial variation in labels (Rolnick et al., 2018). Specifically, we coded a 1D eight-layer fully convolutional neural network (CNN) comprised of seven convolutional and one fully connected layer (size 150) in TensorFlow (Python). We used a stride of four to reduce input size four-fold in each layer. This size reduction dictated the number of layers in the network to support our largest model of 65,536 inputs (2.6s). We used a kernel size of 30, which indicates how many data points each layers sees in the convolution step. Features are built based on the kernel size at each layer. Layers contained the following number of filters: layer 1 had 320, layer 2 had 384, layer 3 had 448, layer 4-7 had 512. The output of the model is 3 values, a probability distribution over wake, non-rem and rem. This model architecture was the same for all input sizes from 1 sample to 2.6 s (65,536 data points). L2 weight regularization with 1e-6 was utilized. Learning rates were progressively reduced from 1e-4 to 5e-6 throughout training. Each layer utilized a standard ReLU activation function with the exception of the final convolutional layer which had no activation function applied. The loss function minimized by the CNN was softmax cross entropy (Good, 1952). CNN confidence was quantified as the entropy of the output distribution scaled to [0, 1]. Model parameters were chosen through a manual process of hyperparameter tuning. The manual hyperparameter tuning follows a coordinate descent approach in which each hyperparameter is varied until an optimal value is identified for that parameter, then the next parameter is varied, holding all others fixed. Hyperparameter selection was conducted early in the experimentation cycle and remained fixed throughout the experiments so as not to introduce added confounds.

In all experiments, the CNN was presented with raw neural data. Two exceptions exist: 1) if data was low-passed below 16 Hz to examine the role of canonical oscillations, or 2) if data was high-passed above 750 Hz to examine the role of high-frequency (predominantly neuronal) activity. When these exceptions occur, it is for the purpose of interpretation and is directly specified in text, figure legends, and often the figure itself. Models were tasked with learning to predict sleep and wake states (REM, NREM, and wake) based on labels generated by a consensus amongst three expert human scorers. In some experiments, raw neural data were progressively ablated (see below) in preprocessing. To avoid common numerical problems, input data were linearly scaled by 10^-3^. The batch size during training was one, and the model was trained for 175,000 training steps. We selected a train/test split such that the training was composed of 18 consecutive hours of data, and the held-out test set comprised 6 h spanning a light/dark transition. Unless otherwise noted, step size was 1/15th of a second to match video fps. Model performance was evaluated using balanced accuracy (Brodersen et al., 2010; Kelleher et al., 2015) unless otherwise noted below. Model code will be available at github.com/hengenlab at the time of publication.

CNN models functioned as follows. At each time step, a model was presented with an interval of data (the duration of this interval was experimentally varied, see below). The model’s task was to assign a probability of REM, NREM and wake to the central sample point. As a result, the task of each CNN was to learn the instantaneous state label at the center of a window of data, with only the information surrounding the center. Crucially, CNNs have no memory, and thus each observed segment of data is an independent decision.

A more simple fully convolutional model was chosen over a more complex model such as ResNet with the following rationale: 1) to reduce computational burden, enabling the training of many thousands of models with varying input conditions, 2) it’s not clear that more complex models benchmarked against unrelated datasets would be a better fit for this data, 3) the existence of substantial label error dictates that there is more value in focusing on data than on model tuning.

#### Basic Accuracy of Sleep and Wake States by Brain Region (Fig. 1D-F)

To address the question of whether sleep and wake states are robustly embedded in the dynamics of the ten brain regions sampled in our recordings, we asked if CNNs could learn to decode sleep and wake from the broadband neural data within a single brain region (64 channels of data). Model input was 2.6 s (65,536 sample points) of 64 channels of raw neural data from each implant site (n = 45 implants from N = 9 animals, 45 total models).

#### Extended Accuracy Evaluation in High Gamma Range (Fig. 1G-H)

To examine the decodability of state information within the high gamma range, we implemented per-brain region models on neural data from 2 distinct animals (n = 2 animals, with 12 probes in total derived from animals 2 and 7). The data had been manipulated via bandpass filtering to confine it to progressively amplified frequency information. Model input consisted of data that underwent the following bandpass filters: broadband, low-pass 16 Hz, 100-200 Hz bandpass, 200-300 Hz bandpass, 300-400 Hz bandpass, 400-500 Hz bandpass. Six models were executed per probe (72 total models). Note that EMG signal can be extracted from 300-500 Hz (Watson et al., 2016). The failure of models to learn in these bands suggests the signal is insufficient to support state classification in the recordings. We further confirmed that in a true EMG recording (750 Hz high-passed) models failed to train. The impact on the models’ capacity to learn from these frequency-specific datasets was closely monitored. Accuracy was evaluated using balanced accuracy directly on CNN test output.

#### Progressive High Pass (Fig. 2A)

To test whether sleep and wake states could be extracted from neural data absent the canonical waves that human scorers rely on, we progressively eliminated slow components of neural dynamics from all implants within an animal and measured the impact on the models’ ability to learn. Model input was ∼1 s (24,576 sample points) of high-pass filtered neural data (3rd order Butterworth) from all channels within an animal (n = 1 animal). 24 models were run. Specifically, models were trained and tested on neural data after the following high-pass filters were applied: 0 (raw), 0.5 Hz, 1 Hz, 2 Hz, 4 Hz, 6 Hz, 8 Hz, 12 Hz, 20 Hz, 50 Hz, 100 Hz, 250 Hz, 500 Hz, 750 Hz, 1,000 Hz, 1,500 Hz, 1,600 Hz, 1,700 Hz, 1,900 Hz, 2,000 Hz, 3,000 Hz, 5,000 Hz, 7000 Hz, 10,000 Hz. Accuracy was evaluated using balanced accuracy directly on CNN test output.

#### Exploration of Inter-Regional Differences in Low Pass, High Pass, and Broadband Data (Fig. 2B)

In order to assess the potential disparities between brain regions apparent in low pass, high pass, and unaltered broadband variations of the data, we implemented an extensive suite of models corresponding to each probe across all subjects (n = 9 mice incorporating 45 probes in total). The identical model structure was applied in all scenarios, wherein the unmodified broadband data was subjected to preprocessing via a low pass filter at 16 Hz, a high pass filter at 750 Hz, or was left unfiltered. For each model, a single probe targeting a specific brain region (comprising 64 channels) was utilized, with a data input duration of 2.6 seconds (encompassing 65,536 data points). Accuracy was evaluated using balanced accuracy directly on CNN test output.

#### Incremental Diminution of Input Size (Fig. 2C-D)

In pursuit of ascertaining the minimal temporal duration required to accurately decode sleep and wake states, we orchestrated an expansive set of models corresponding to each probe in all subjects (n = 9 mice comprising 45 probes in total), and trained them on an input size that was systematically reduced. Each model was informed by data from a single implant located within a particular brain region. A model with identical dimensions and hyperparameters was employed in each training phase, with the variation of increasingly truncated input sizes. The model’s task was to generate predictions solely based on the temporal window of data presented, devoid of any prior knowledge pertaining to the animal’s state. The progressive series of input sizes utilized were as follows: 2.6 s (65,536 data points), 1.3 s (32,768 data points), 327 ms (8,192 data points), 82 ms (2,048 data points), 41 ms (1,024 data points), 10 ms (256 data points), 5 ms (128 data points), 2.5 ms (64 data points), 1.3 ms (32 data points), 0.6 ms (16 data points), 0.3 ms (8 data points), 0.16 ms (4 data points), 0.08 ms (2 data points), and 0.04 ms (1 data point). Accuracy was evaluated using balanced accuracy directly on CNN test output.

#### Model Training and Testing on Low-Pass Filtered Data (Fig. 3D)

Data for this experiment were prepared using two manually selected channels, both spiking and non-spiking, with clear local field potentials. In some cases only one channel was included due to simple computational reasons. No exclusion criteria was applied beyond selecting only high quality channels. Each channel’s data underwent two transformations: in the first condition, it was left in its original, temporally intact form; in the second, the data was shuffled, reordering the data within each sampled segment. The shuffling process was executed by randomly drawing samples from the dataset and shuffling the data points within the sample, maintaining the same temporally intact sampling method as used for the unshuffled data. The models were then trained on the transformed data using a variety of input sizes from 2.6 seconds (65,536 data points) to 0.04 milliseconds (1 data point), generating a total of 2,028 single-channel models. Model performance was evaluated using balanced accuracy for each scenario.

#### Model Training and Testing on High-Pass Filtered Data (Fig. 3E)

The same channels and range of input sizes as in Fig.3D were used for this part of the experiment. However, the preprocessing stage was different, employing a high-pass filter set at 750 Hz to remove low-frequency details from the raw data. The models were then trained on both the unaltered and shuffled high-pass data.

#### Evaluation of Model Accuracy vs. Input Size (Fig. 3F)

The data displayed in 3F is the same data from 3D and 3E, presented as a function of the input size.

### Flicker Experiments

#### Flicker definition (Extended Data Fig. 7)

We applied a series of criteria to CNN output to identify flickers for inclusion in our analyses. To start, we trained three CNNs to identify NREM, REM and wake based on raw broadband data (1 s input) from all 64 channels contained in each implantation site (n = 45 implantation sites, 9 animals, 126 models). As a result, we had triplicate CNN-generated state scores for each recorded brain region. This was to avoid sporadic random error due to subtle inconsistencies in training. Often, we observed that CNN output would preemptively begin to increase confidence in an incoming state a few seconds prior to a global transition. To avoid transition-related ambiguity, we generally did not consider flickers within 30 s of a global transitions, but to account for the fact that some transitions were modified in time (e.g., a slow transition from quiet wake to NREM sleep). We manually evaluated CNN confidence surrounding each global state transition in the 24h of data from each implantation site in each animal in each replicate of the CNN (n = 45 implantations from 9 animals, 3 replicates). We extended the window of exclusion around transitions in which evidence of the incoming state was present beyond the 30 s window. We then applied a series of confidence filters to the remaining data in each replicate. To avoid general periods of low confidence output, we identified and excluded any 35 s epochs with a mean confidence <75%. To restrict our analyses to high confidence flickering, we next eliminated 1s epochs with a mean confidence <75%. Together, these criteria excluded 20.91% of our recordings. In each replicate, we then assigned the high confidence state label to each time step (1/15th of a second), and collapsed the three model outputs into a single array by selecting the majority state at each interval. We then slid a 35 s rolling mode filter across the majority state array to create a label corresponding to the stable macro state surrounding every point in time. We defined flickers as disagreements between these two arrays.

We had two further exclusion criteria to avoid overlap between flickers and previously described episodic arousal phenomena. First, flickers which co-occured across all probes in the animals were excluded. Specifically, this includes microstates, such as microsleeps and microarousals. Due to the 35 s modal filter of the majority state array, these had a maximum duration of ∼20 seconds. Second, please note that flickers were detected in raw broadband data. However, to avoid overlap with events that could be visible in low-frequency data, we also detected flickers in models trained on data low-passed below 16 Hz. We excluded any flickers detected in the broadband which had any temporal overlap with a flicker detected in any region of the low-pass. This excludes low frequency phenomena previously described in EEG & LFP such as local slow waves in wake and REM.

We also observed high-confidence, localized CNN errors (similar to flickers) amidst transitions between states (Fig. 6C). Consistent with progression along a transition’s time course, we found their rate of occurrence was inversely correlated with time to/from a human-labeled state change (p=0.016892). Because they have a similar confidence profile in the CNN’s predictions to flickers and they reflect transition dynamics, we used these as samples of transition spiking activity. To extract these events, we performed a comparable procedure to the previously mentioned flicker detection with two exceptions: 1) rather than excluding intervals of transitions, we excluded the complement (all non-transition intervals), 2) we did not exclude general periods of low confidence (based on 35s window) because the time course of a transition is often a low confidence period.

#### Synthetic Flickers (Fig. 5C)

To quantify the ability of the CNN to accurately detect brief intervals of a state **B** embedded within a containing state **A**, we constructed synthetic flickers. To do this, we identified all intervals of state **A** (NREM, REM, and wake) that were assigned the same label by all three human scorers, as well as the standard 2.6s CNN model (confidence >90%. See: Basic Accuracy of Sleep and Wake States by Brain Region). High confidence intervals were a minimum of 3 s in length. To simulate state **B**, we spliced segments of each state **A** into each other state (6 total combinations of REM, NREM, and wake into one another). Splices were 9.5, 19.0, 28.6, 47.6, 66.7, 133.3, and 333.3 ms. 100 splices of each duration were randomly selected from all high confidence intervals in the 24 h period and pasted into a randomly selected high confidence interval of another state. To illustrate with a specific example, consider a continuous segment of 47.6 ms of high confidence REM. This was randomly selected from thousands of >3 s intervals. A high confidence segment of wake was chosen at random from thousands of examples, and the 47.6 ms REM splice was pasted into a random location in the selected segment of wake. The insertion of the REM splice overwrote the corresponding section of wake data. We then asked whether splices were correctly identified by a CNN with a 655 ms window and a step size of 9.5 ms. We chose this step size to establish a functional lower limit of sensitivity while maintaining a reasonable computational load. We evaluated whether the CNN correctly identified any portion of the spliced state **B**.

#### Coflicker Analysis (Fig. 5F)

To understand whether flickers are a locally regulated (independent) or global phenomenon, we calculated the conditional probability of all pairs of regions flickering simultaneously according to:

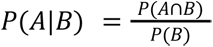

#### Single Unit Analysis (Fig. 6D,E,F)

Principal Component Analysis (PCA) was employed on each 12-hour spike-sorted block of CNN-classified regional recordings to evaluate whether single unit spiking could reliably discriminate arousal states. For each arousal state (wake, NREM, and REM) and flicker/transition type state (wake-to-nrem, wake-to-rem, nrem-to-wake, nrem-to-rem, rem-to-wake, and rem-to-nrem) we aggregated all interspike intervals (ISIs) for each neuron present during that state. We then randomly sampled 100 ISIs from these aggregated ISIs in contiguous runs of 1-10 ISIs (short snippets). We repeated this 100 times for each state. For each sample of a neuron’s spiking we calculated the mean ISI of the neurons (the inverse-mean instantaneous firing rate-gave similar results for this analysis). Prior to PCA, for each flicker type, we z-score normalized the firing rates for each neuron in the performed PCA. This z-score normalization was fit to the samples from the surrounding state and the predicted state (e.g. for a wake-to-nrem flicker, Wake and NREM appropriately). PC1 consistently separated these two classes with little overlap. We then transformed samples from the appropriate flicker-type and transition onto this axis. Transitions were available for four of the six state-pair combinations (not wake-to-REM or REM-to-NREM because they are not commonly observed). To enable us to evaluate trends across clustering blocks, we MinMax scaled the projections of samples along PC1. PCA projections were multiplied by -1 if the mean value of the projections of the predicted state were less than the mean value of the projections of the surrounding state. This aided visualization by ensuring samples of the surrounding state took on negative values and samples of the predicted state took on positive values to make visualization. To avoid the possibility of overestimating statistical effects, we took the mean projection for each surrounding state, predicted state, flicker state, and transition state in a clustering block. Error bars were challenging to formulate due to ISI resampling and possible dependency between spiking in two regions during co-flickering. A solution which is intuitively similar to standard deviation, with a focus on capturing animal-wise variance, was used. We first grouped all samples of spiking from a clustering block to create a mean for each. The SEM was then calculated with degrees of freedom multiplying the SEM by the square root of the number of animals (i.e. number of uncorrelated observations). This gave error bars that were roughly reflective of the trends seen in independent statistical significance tests. This is intuitively similar to the standard deviation. For each flicker type we fit a linear mixed effects model relating state to PC1 projection with animal as a random effect. We then performed an ANOVA and post-hoc EMMeans with Tukey correction. When grouping flickers by flicker-type we multiplied p-values by 6 for Bonferroni multiple hypothesis correction. When grouping flickers by region we multiplied p-values by 10.

To evaluate the directional perturbation of portions of the population, we used the same sampled data used in PCA. For each flicker type, we iterated through each of the 100 samples for the surrounding state, flicker state, transition state, and predicted state in parallel. For each neuron, for each state we identified whether it or the surrounding state had the higher instantaneous firing rate. For each sample we evaluated the percent of neurons whose firing rate was increased (by any amount) relative to the sample of the surrounding state. As a negative control, we compared intervals of the surrounding state to eachother (sample 1 to sample 100, sample 2 to sample 99,…). For this negative control, the portion of units with increased firing should ideally be ∼50% (representing chance), however in the case of some rare flicker types, it deviated from this due to sampling error.

#### Flickers During Waking Behavior(Extended Data Fig. 4G, Fig.7B,E,H)

To correlate animal behavior with flickers we aligned video (15 fps) with neural data using a dedicated digital data channel synchronized with electrophysiological data acquisition, providing nanosecond-accurate timestamps per video frame (E3Vision, WhiteMatter, LLC, Seattle, WA). We use Farnebäck dense optical flow (Farnebäck, 2003) to measure movement. Specifically, motion was measured using dense optical flow which provides a per-pixel movement vector calculated between temporally adjacent video frames. Dictated by the video sampling rate, 15 per-pixel optical flow vectors were produced per second of recording. A video frame is segmented into three notable parts; 1) the subject animal, which is the core focus, 2) the background which should be excluded, and 3) the headstage tether cable which should be excluded, but produces the highest movement vectors due to its contrast with the background, immediate proximity to the camera, and quick jittery motion. To reduce the effect of cable movement, we excluded the top 3% of motion vectors. To focus analysis on the animal (as opposed to the background) we took the remaining top 10% of motion vectors. Explicitly, this retained the 87^th^ percentile to 97^th^ percentile of motion vectors produced by dense optical flow. Total animal movement was computed as the mean of these motion vectors for each frame.

We next calculated two normalizations to allow comparisons across animals and over time. To accommodate drift and environmental changes, such as light/dark, we rescaled all motion vectors per each hour of video to a [0, 1] range. To normalize differences between mice we computed the 75^th^ percentile of motion vectors per mouse and rescaled the movement values associated with each frame such that each animal’s 75th percentile movement values were aligned. Finally, we collapsed movement values above 1 to 1, resulting in a movement vector in the [0, 1] range. In other words, periods of high movement appeared as a sustained sequence of 1s. These normalization steps produced a movement value that is sufficiently invariant to light/dark cycles, differences in recording resolution, and variations in camera orientation and zoom to align locomotor states across animals.

Three substates of waking were defined. Specifically, sustained high activity, low to intermediate activity, and brief pauses embedded within high activity. We computed two median filters over the normalized per-video frame movement values. Median filters are particularly useful in this context because they produce a smoothing with a sharp contour to the data that is cleanly delineable. The first was a rolling median using a 60 s window. At a threshold of 0.75, this median filter broadly segmented periods of sustained, high activity and periods of intermediate to low activity. Within periods of high activity, we then used a rolling median using a 0.66 s window to segment pauses from within sustained high activity. We manually evaluated a random subset of each locomotor state to confirm the accuracy of the algorithm parameters.

#### Flickers During Sleeping Behavior(Fig.7A,C,D,F,G,I)

Using an analogous process to wake, three substates of sleep were defined. Specifically, sustained high activity (indicative of an extended microarousal), low activity, and brief high activity embedded within low activity (indicative of a twitch).

To capture minor fluctuations in optical flow (indicative of brief stillness, or “freezing”) we inverted the trace of the magnitude of the optical flow and applied the scipy find_peaks function with default hyperparameters. We identified freezing as the 1 second following each of these peaks.

### Software and Statistical Analyses

Data are reported as mean ± SEM unless otherwise noted. We used a mixed effects regression analysis with animal as a random effect followed by EMMeans with Tukey post hoc correction (LME4, R) (Bates et al., 2015) to determine statistical significance (p<0.05) unless otherwise noted. Mixed effects models comparing accuracy of models considered only models which trained successfully (accuracy > 33%).

### Data Availability

The datasets generated and/or analyzed in this study constitute tens of terabytes of raw neural broadband. The data are stored in a cost efficient manner not immediately accessible to the internet. We are excited to share data upon reasonable request, and as technical limitations make possible.

### Code Availability

All relevant code from our lab, including software needed to run recordings or CNN models like ours is in python3 and will be publicly available at https://github.com/hengenlab at the time of publication. Other groups’ code including Open Ephys, SpikeInterface, and MountainSort4 is publicly available as specified in Methods.

## Acknowledgments

This work is supported by NIH BRAIN Initiative 1R01NS118442-01 (KBH), and the Schmidt Futures Foundation SF 857 (D.H.). Through the Pacific Research Platform, this work was supported in part by NSF awards CNS-1730158, ACI-1540112, ACI-1541349, OAC-1826967, the University of California Office of the President, and the University of California San Diego’s California Institute for Telecommunications and Information Technology/Qualcomm Institute. Thanks to CENIC for the 100 Gpbs networks.

## Author Contributions

D.F.P. developed and ran the models and performed behavioral analyses, A.M.S. performed statistical analyses, wrote the paper, performed the single unit spiking analyses, and performed behavioral analyses, Y.X. provided sleep scoring expertise and animal care, S.J.B contributed substate analyses, S.F. provided sleep scoring expertise, D.T. performed flicker identification, T.B. provided intellectual and technical consultation, E.L.D. provided mentorship and consultation, D.H. provided mentorship and consultation, K.B.H. led, directed, and envisioned the project, edited figures, and wrote the paper.

## Materials & Correspondence

All correspondence should be directed to Keith Hengen at

